# Premyelinating Oligodendrocyte Survival Governs CNS Remyelination

**DOI:** 10.64898/2026.02.23.707571

**Authors:** Michael E. Stockton, Maureen Wentling, Yessenia I. Mancha Corchado, Abigail Kucera, Stephanie Karas, Tania A. Seabrook, Clara Mutschler, Lindsay A. Osso, Kristen H. Schuster, Gustavo Della-Flora Nunes, Anthony R. Chavez, Ethan G. Hughes

**Affiliations:** Dept. of Cell and Developmental Biology, University of Colorado School of Medicine, Aurora, CO, USA; Myrobalan Therapeutics, Inc., Medford, MA, USA

## Abstract

Myelinating oligodendrocytes are produced throughout life by the constitutive differentiation of oligodendrocyte precursor cells (OPCs). However, the rate of oligodendrocyte generation changes with age and after a demyelinating injury. Here, we report that variation in premyelinating oligodendrocyte (preOL) survival modulates the rate of oligodendrocyte production. PreOL survival increased to drive the regeneration of oligodendrocytes after demyelination and decreased in middle-aged mice to contribute to age-related decline in oligodendrocyte production. Furthermore, we demonstrate that treatment with a GPR17 antagonist, Myro-02, increases oligodendrocyte replacement by promoting preOL survival after demyelination. Together, our findings demonstrate that increased survival of preOLs governs the regeneration of oligodendrocytes following demyelinating injury and suggest that modulating preOL survival may be an alternative therapeutic avenue to promote oligodendrocyte regeneration.

---

Myelin sheaths generated by oligodendrocytes accelerate action potential propagation and provide essential metabolic support to axons in the central nervous system (CNS). Throughout life, new oligodendrocytes arise from oligodendrocyte precursor cells (OPCs) via a process called oligodendrogenesis^1–3^. OPCs exit the cell cycle and differentiate into premyelinating oligodendrocytes (preOLs), a transient cellular stage immediately prior to myelin formation^4,5^. PreOLs halt cellular migration and filopodial activity, and extend long, slender processes that do not yet enwrap axons^6,7^. PreOLs are poised to make critical fate decisions between cell death and survival^8^, and are highly vulnerable to apoptosis^7,9,10^. Indeed, genetically blocking apoptosis in the oligodendrocyte lineage during development results in ectopic production of oligodendrocytes and myelin^11,12^. These observations raise the possibility that the preOL stage serves as a critical decision point that determines how many new oligodendrocytes ultimately integrate into neural circuits.

Demyelinating injuries typically provoke a regenerative response that produces a surge of new oligodendrocytes and remyelination^13–18^ with parenchymal OPCs giving rise to the majority of these new oligodendrocytes^19–21^. In human multiple sclerosis (MS) lesions, OPCs are commonly present^22,23^, and can differentiate into preOLs^17,24^. Yet, remyelination remains variable and is often incomplete ^25,26^, suggesting that there is additional failure of either OPC differentiation or preOLs survival. Recent work shows that the rate of OPC differentiation is not increased after demyelinating injury^27^, suggesting additional downstream checkpoints may govern remyelination.

Aging is associated with reduced remyelination efficiency^28,29^, likely arising from convergent impairments in OPC proliferation^30,31^, differentiation responsiveness^27,28,32^, and potentially preOL survival^7,9,10^. Yet even in the aging brain, measurements of preOL survival have not been performed making its contribution to the failure of oligodendrocyte regeneration unclear. Understanding how these cellular decisions change across the lifespan will be essential for interpreting remyelination failure in older individuals where diseases, such as multiple sclerosis, are most severe.

Therapeutic strategies for remyelination have largely focused on enhancing OPC differentiation, from early work on thyroid hormone, glucocorticoids, and retinoids^33–35^ to more recent high-throughput identification of compounds like clemastine^15,36,37^. However, these “remyelination therapies” have had limited success in clinical trials^38^. Since a substantial fraction of differentiation attempts are lost at the preOL stage in the healthy adult cortex^10^, increasing OPC differentiation may be insufficient to promote regeneration unless coupled with improved preOL survival and integration. Therefore, promoting preOL survival represents an alternative and largely unexplored therapeutic axis. Recently, GPR17, a G-protein coupled receptor enriched in preOLs^4,5^, has been proposed to regulate oligodendrocyte maturation^39^ and genetic studies, as well as pharmacological inhibition of GPR17, suggest that blocking this pathway can enhance remyelination^40,41^. These findings raise the possibility that GPR17 antagonism could enhance repair by improving the survival of differentiating preOLs.

Here, we performed quantitative lineage tracing via longitudinal *in vivo* two-photon imaging in the mouse cortex to define the role of preOL survival in oligodendrocyte generation during homeostasis, demyelination, and repair in the young adult and middle-aged brain. This approach allows us to dissect, with temporal precision, how often OPCs proliferate, differentiate, how frequently preOLs survive, and how these events contribute to oligodendrocyte generation throughout adulthood. We find that across conditions the rate at which OPCs initiate differentiation remains constant, but preOL survival is dynamically regulated and emerges as the key determinant of regenerative success. In middle-aged mice, although OPCs preserve the ability to replace lost oligodendrocytes, we find that failure to recruit dormant OPCs, which did not proliferate or differentiate over 12 weeks, and diminished preOL survival of non-dormant OPCs together limit remyelination. Finally, pharmacological antagonism of GPR17 with Myro-02 in young adult mice selectively increases preOL survival leading to the enhanced replacement of lost oligodendrocytes, without measurably changing OPC proliferation or differentiation. Together, these findings establish preOL survival as a central, tunable determinant of oligodendrocyte regeneration efficiency and suggest that modulating premyelinating oligodendrocyte survival may be an alternative therapeutic avenue to promote remyelination.

## Results

### Premyelinating oligodendrocyte survival is inefficient in the healthy adult cortex

In the adult brain, the majority of premyelinating oligodendrocytes are thought to fail to integrate as mature, myelinating oligodendrocytes ^10^. To monitor the differentiation of OPCs and determine the fate of preOLs in the intact adult brain, we developed a quantitative lineage tracing approach. To do this, we generated triple transgenic mice (*Olig2-CreER;R26-lsl-tdTomato;Mobp-EGFP)* wherein tdTomato is expressed across the entire oligodendrocyte lineage but EGFP expression is limited only to mature oligodendrocytes and their myelin sheaths (**Figure 1A and 1B**). Importantly, EGFP expression driven by the *Mobp* promoter only occurs following the successful integration of a myelinating oligodendrocyte and the formation of myelin sheaths.

**Figure 1.**
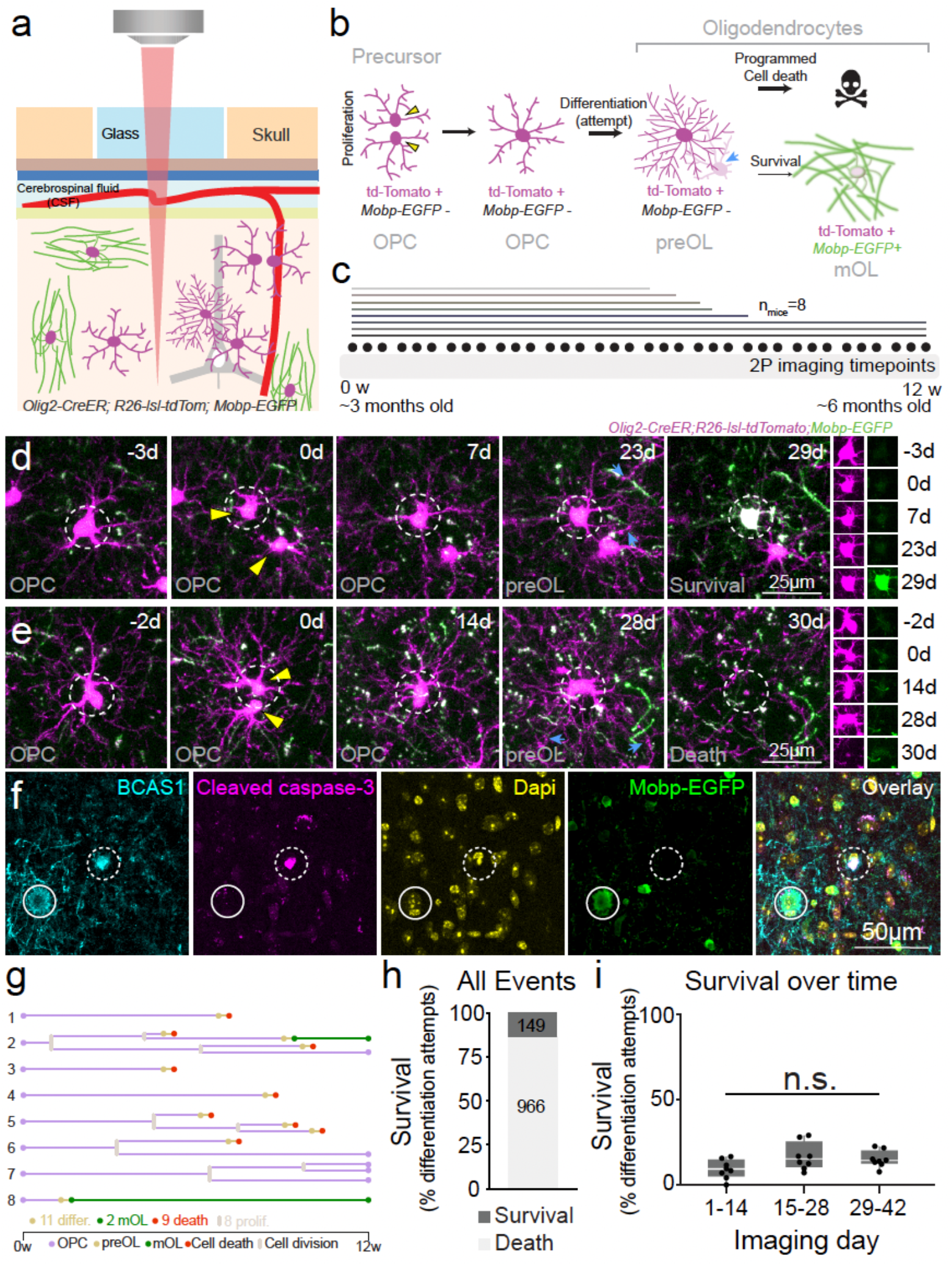
Quantitative lineage tracing reveals premyelinating oligodendrocyte survival is inefficient in adult mice. **a**, Schematic of *in vivo* two-photon imaging of *Olig2-CreER; R26-lsl-tdTomato;Mobp-EGFP* mice. **b**, Cartoon of cellular behaviors in the oligodendrocyte lineage (oligodendrocyte precursor cell, OPC; premyelinating oligodendrocyte, preOL; myelinating oligodendrocyte, mOL). Blue arrow indicates territory invasion of neighboring OPCs. **c**, Experimental design, lines represent the imaging duration for individual mice (n_mice_=8). **d**,**e**, Representative images of an OPC undergoing cell division followed by OPC differentiation into a preOL. The preOL then survives and generates mOL (**d**) or undergoes cell death (**e**). Yellow arrowheads= cell division. Blue arrows= territory invasion by neighboring OPC. Far right column shows tdTomato and EGFP signal in soma for cell of interest at each timepoint. **f**, Representative images showing an apoptotic preOL (Dashed circle: fragmented BCAS1^high^ *Mobp*-EGFP+; cleaved caspase-3+) and a newly generated mOL(Solid circle: BCAS1^high^ *Mobp*-EGFP+; cleaved caspase-3-). **g**, Representative quantitative lineage tracing diagrams of individual OPC clones. OPC differentiation attempt (yellow), proliferation (grey), cell death (red), and new mOL generation (green). **h**, Percentage of differentiation events that result new mOL generation across the entire imaging duration. n_mice_ = 8, n_events_ = 1115 (84-191 events/animal). **i**, Quantification of percentage of differentiation events that result in new mOL generation over time. H(2) = 5.0381, p=0.0805, Kruskal-Wallis Test. n_mice_ = 8. Dots represent animal means. Box and whisker plots denote median and IQR. For all analyses: n.s. not significant, *p < 0.05, **p < 0.01, ***p < 0.001. For detailed statistical information see Supplementary Table 1.

Consistent with the long-lived nature of myelinating oligodendrocytes and their associated myelin sheaths in healthy adult mice^42^ and in humans^43^, *Mobp*-EGFP-positive oligodendrocytes remain stable across many months^10^.

First, we assessed the penetrance of *Olig2* promoter driven tdTomato expression across the oligodendrocyte lineage in young adult triple transgenic mice (3-6 month old). We found that tamoxifen-induced Cre recombination resulted in tdTomato expression in 98.21 ± 0.89% of PDGFRa+ oligodendrocyte precursors and in 91.58 ± 1.72% of ASPA+ mature oligodendrocytes in the motor cortex (Layers 1-6), indicating this approach reliably labeled nearly all oligodendrocytes and their precursors. Next, to examine oligodendroglia behaviors in the intact brain, we implanted cranial windows over the motor cortex and used long-term *in vivo* imaging to acquire the same field of view three times per week for up to 12 weeks (**Figure 1A-C**). To monitor OPC proliferation, differentiation attempts, and the fate of preOLs over time, we used quantitative linage tracing. To do this, we reconstructed individual cell lineage trees for over 1400 OPC clones from eight healthy young adult animals and assessed cellular behaviors with observation periods between seven and twelve weeks (**Figure 1C**). Oligodendrocyte precursor cells (OPCs= tdTomato+, *Mobp*-EGFP-) exhibited dynamic processes with motile filopodia and underwent cellular division (**Figure 1D and 1E, Supplementary Video 1**). OPCs were present throughout the parenchyma, displaying robust homotypic repulsion that maintained exclusive cellular territories from neighboring OPCs (**Figure S1A, Supplementary Video 2**) as previous described^6^. As OPCs initiated differentiation, they ceased both migration and filopodial activity, instead developing long, slender processes typical of premyelinating oligodendrocytes. This transition was marked by the encroachment of neighboring OPC processes into the differentiating cell’s territory and the expression of BCAS1 (**Figures 1D-E and S1B-E**). Premyelinating oligodendrocytes (preOLs= tdTomato+, *Mobp*-EGFP-) exhibited increased branching, branch length, and increased total cell size compared to OPCs (**Figure S1F-K**). PreOLs that successfully integrated into the cortical environment as myelinating oligodendrocytes (mOL= tdTomato+, *Mobp*-EGFP+) retained tdTomato expression throughout the imaging period and expressed *Mobp*-EGFP following the generation of myelin sheaths (**Figure 1D**; **Supplementary Video 3**). TdTomato-positive preOLs that failed the differentiation process and did not survive exhibited cellular fragmentation *in vivo*, and were BCAS1-positive, *Mobp*-EGFP-negative, and cleaved-caspase 3-positive in histologically assessed tissue (**Figure 1F, Supplementary Video 4**), indicating these preOLs underwent programmed cell death. We applied quantitative linage tracing (**Figure 1G**) to over a 1000 OPC differentiation attempts across eight mice and found that only 13.36% of preOLs survived to generate myelinating oligodendrocytes (149 / 1115 cells; n = 8 mice, aged 3–6 month; **Figure 1H**). Finally, we did not observe any direct death of OPCs in this mouse line. Every OPC disappearance was preceded by the generation of a preOL, which is consistent with recent work that showed PDGFRα-positive OPCs are not labeled with cleaved-caspase-3^16^

Oligodendrocyte precursors undergo constant proliferation and differentiation to maintain their population and generate new myelinating oligodendrocytes in the adult brain^6^. To examine if there were time-dependent changes in these oligodendroglia behaviors, we analyzed 14-day temporal bins for six weeks. Similar to previous studies^14,18,44^, we found that the number of new oligodendrocytes generated does not vary across time in the young adult cortex (*P =* 0.3847, F(2,23) = 1.0, One-way ANOVA, **Figure S2A**). We found that the number of OPCs that attempted differentiation or underwent cell division was not altered over the 42 day imaging period (Differentiation: *P =* 0.1631, F(2,23) = 1.98, One-way ANOVA. Proliferation: *P =* 0.7939, F(2,23) = 0.23, One-way ANOVA. **Figure S2B and S2C**). We assessed the proportion of preOLs that survived the differentiation process within each temporal bin and did not observe any change in preOL survival over the 42 day imaging period indicating that preOL survival is constant in the healthy young adult cortex (*P* = 0.0805, H(2) = 5.0381, Kruskal-Wallis Test, **Figure 1I**).

### Quantitative lineage tracing permits determination of oligodendrocyte loss and regeneration

The cellular principles that govern regenerative oligodendrogenesis and remyelination remain unclear. To examine oligodendroglia lineage progression following demyelinating injury, we implemented our quantitative lineage tracing approach. We elicited myelin loss and oligodendrocyte death by treating young adult (3-4 month old) *Olig2-CreER;R26-lsl-tdTomato;Mobp-EGFP* mice with a 0.2% cuprizone diet (**Figure 2A**). Chronic cranial windows were implanted over the motor cortex and long-term two-photon *in vivo* imaging was used to acquire the same field of view three times per week in mice before, during, and after cuprizone administration (**Figure 2B**). A cohort of healthy mice followed the same experimental design but were administered a diet without cuprizone (**Figure 1**). We monitored oligodendrocyte loss and regeneration while simultaneously reconstructing cell lineage trees of individual OPC clones (Healthy, n_mice_ = 8; Cuprizone, n_mice_ = 9). Consistent with previous findings, we observed no loss of oligodendrocytes in healthy mice whereas we found that 53.78-93.46% of pre-existing mature oligodendrocytes were lost in mice fed a 0.2% cuprizone diet for three weeks (**Figure S2D and 2C**). Despite this regenerative response, the population of myelinating oligodendrocytes did not fully recover to healthy or pretreatment levels across seven weeks following removal of cuprizone treatment (**Figure 2D**). Following the removal of the cuprizone diet, we found a robust regenerative oligodendrogenesis response via longitudinal two-photon imaging that was proportional to the amount of oligodendrocyte loss (Spearman’s ρ = 0.88, *P* < 0.0001) that was not observed in age-matched mice fed a normal diet (**Figure S2D and 2C**).

**Figure 2.**
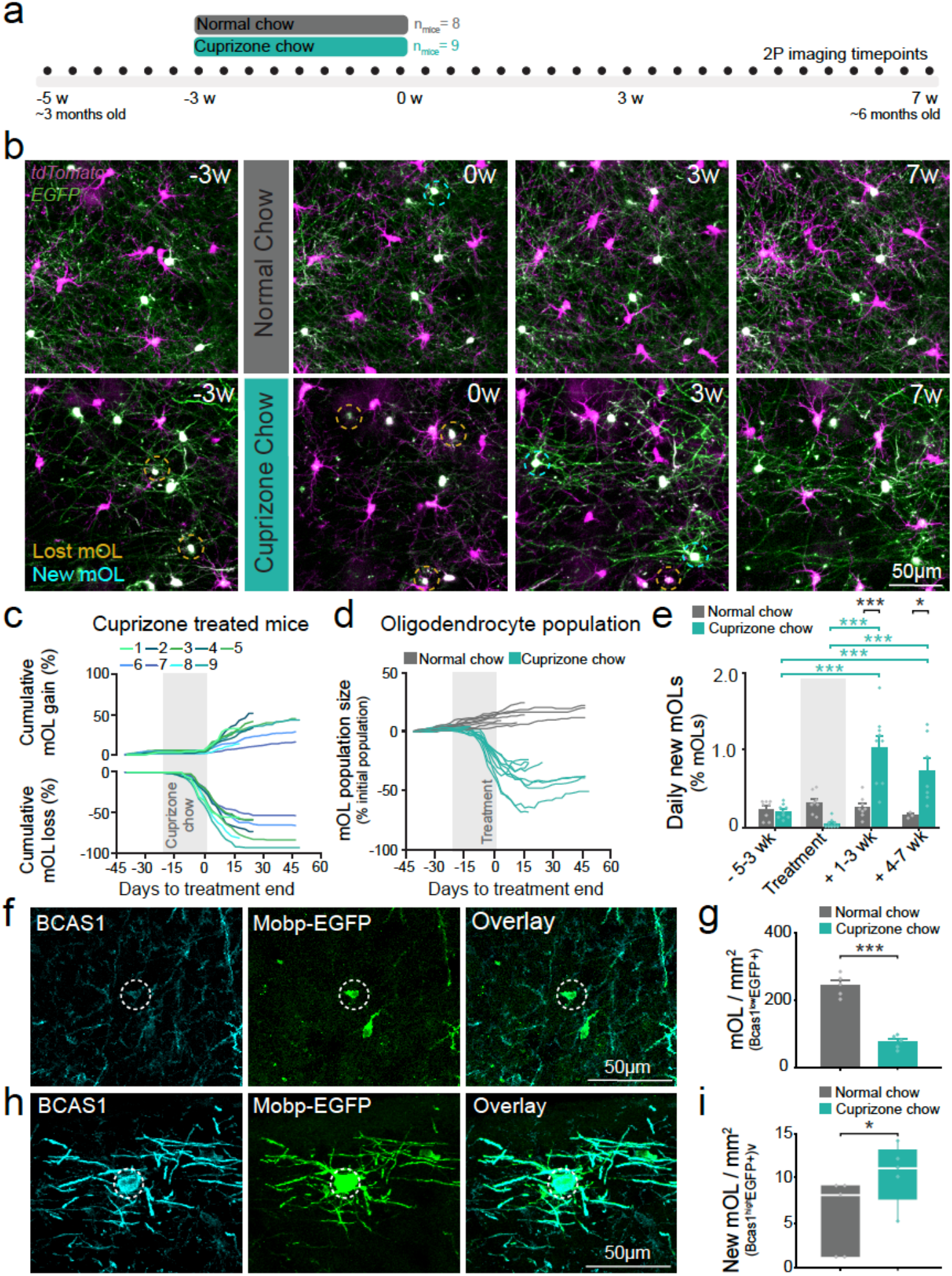
Quantitative lineage tracing permits determination of oligodendrocyte loss and regeneration. **a**, Experimental design for *in vivo* imaging across demyelination and repair in young adult mice. Healthy control mice are the same animals used in Figure 1 (n_healthy_ = 8, n_cuprizone_ = 9). **b**, Representative images showing OPCs and myelinating oligodendrocytes in mice fed a normal diet (top) or a 0.2% cuprizone diet (bottom). Cyan dashed circle: new oligodendrocyte. Orange dashed circle: lost oligodendrocyte. **c**, Cumulative oligodendrocyte gain and loss as a percentage relative to the oligodendrocyte population on day one of imaging in cuprizone mice. Color coded lines represent individual mice. Grey box= cuprizone treatment. **d**, Oligodendrocyte population size relative to day one of imaging in healthy and cuprizone fed mice. Lines represent individual mice and are color coded based on treatment. Grey box= cuprizone treatment. **e**, Quantification of the daily rate of oligodendrocyte generation. There was a significant interaction between treatment and phase, F(3, 40.23) = 21.29, p < 0.0001. Post-hoc comparisons with Tukey’s HSD.. **f**, Representative image from an *Mobp*-EGFP mouse showing a BCAS1^low^*Mobp*-EGFP+ myelinating oligodendrocyte. White dashed circle denotes cell of interest. **g**, Quantification of BCAS1^low^*Mobp*-EGFP+ myelinating oligodendrocytes mice 4 days post-cuprizone versus healthy controls. n_healthy_ = 5, n_cuprizone_ = 5. Pooled t-test, t(8) = 9.16, p<0.0001. Dots represent individual mice. Plots denote mean and S.E.M. **h**, Representative image from an *Mobp*-EGFP mouse showing a BCAS1^High^*Mobp*-EGFP+ new myelinating oligodendrocyte. White dashed circle denotes cell of interest. **i**, Quantification of BCAS1^High^*Mobp*-EGFP+ new myelinating oligodendrocytes 4 days post-cuprizone versus healthy controls. n_healthy_ = 5, n_cuprizone_ = 5. Wilcoxon exact test, p=0.0476. Dots represent individual mice. Box plots represent median and IQR. Error bars represent S.E.M. Dots denote animal means. Error bars represent S.E.M. Dots denote animal means. For all analyses: n.s. not significant, *p < 0.05, **p < 0.01, ***p < 0.001. For detailed statistical information see Supplementary Table 1.

We used quantitative linage tracing to compare new oligodendrocyte generation across different phases of demyelination and repair as well as in relation to healthy control mice. Consistent with previous findings^14,15,18^, we found that cuprizone treatment modulated the generation of new oligodendrocytes. Specifically, oligodendrocyte generation was increased compared to healthy mice during the first three weeks after treatment and from weeks four to seven after treatment (REML mixed model, Tukey’s HSD test, + 1-3wk: *P* < 0.0001, +4-7wk: *P* = 0.0308, **Figure 2E**). Moreover, the daily rate of oligodendrocyte generation within the group of demyelinated mice was increased during the first three weeks after cuprizone compared to pre-treatment and during treatment; an effect that lasted until seven weeks after cuprizone (REML mixed model, Tukey’s HSD test, **Figure 2E**; **S3A**). We corroborated these findings via histology, the density of mature myelinating oligodendrocytes (BCAS1^low^, *Mobp*-EGFP+) was reduced four days after the removal of cuprizone treatment compared to healthy age-matched controls (Healthy: 279.00 ± 17.57 vs. 4d post-cuprizone: 86.60 ± 11.50, *P* < 0.0001, t-test; **Figure 2F and 2G**). Furthermore, we found an increased density of newly generated myelinating oligodendrocytes (BCAS1^high^, *Mobp*-EGFP+) on the fourth day after removal of the cuprizone diet (Healthy: 5.60 ± 1.89. 4d post-cuprizone: 10.40 ± 1.50, *P* = 0.0476, Wilcoxon Exact test, **Figure 2H and 2I**).

Production of new oligodendrocytes via asymmetric division is rare in health and during regeneration

Previous work shows that OPCs that undergo cell division can give rise to oligodendrocytes after a demyelinating injury^45^. Furthermore, after a demyelinating injury in development, there is an increase in the percentage of OPC divisions with an asymmetric fate, where daughter cells consist of an OPC and an oligodendrocyte ^46^. Therefore, it is possible that enhanced OPC proliferation together with an increased frequency of asymmetric division may contribute to regeneration in the adult brain. To determine if OPC proliferation is modulated across demyelination and repair, we quantified the timing of OPC divisions and found that OPCs continuously undergo cell division in both healthy and demyelinated mice (**Figure 3A**). Cuprizone treatment modulated the proliferation of OPCs over time, with a suppression of proliferation during treatment compared to pre-treatment and the first three weeks after treatment (-5-3wk vs. Treatment: *P* = 0.0136. Treatment vs. +1-3wk: *P* < 0.0001, Tukey’s HSD test, **Figure 3B**; **S3C**), but post-hoc comparisons did not reveal any differences between healthy control mice and cuprizone-treated mice (*P* = 0.0015, F(3,40.07) = 6.18, REML mixed model, **Figure 3B**). Thus, increased oligodendrocyte production during repair in cuprizone-treated mice versus controls does not appear to be driven by increased OPC proliferation.

**Figure 3.**
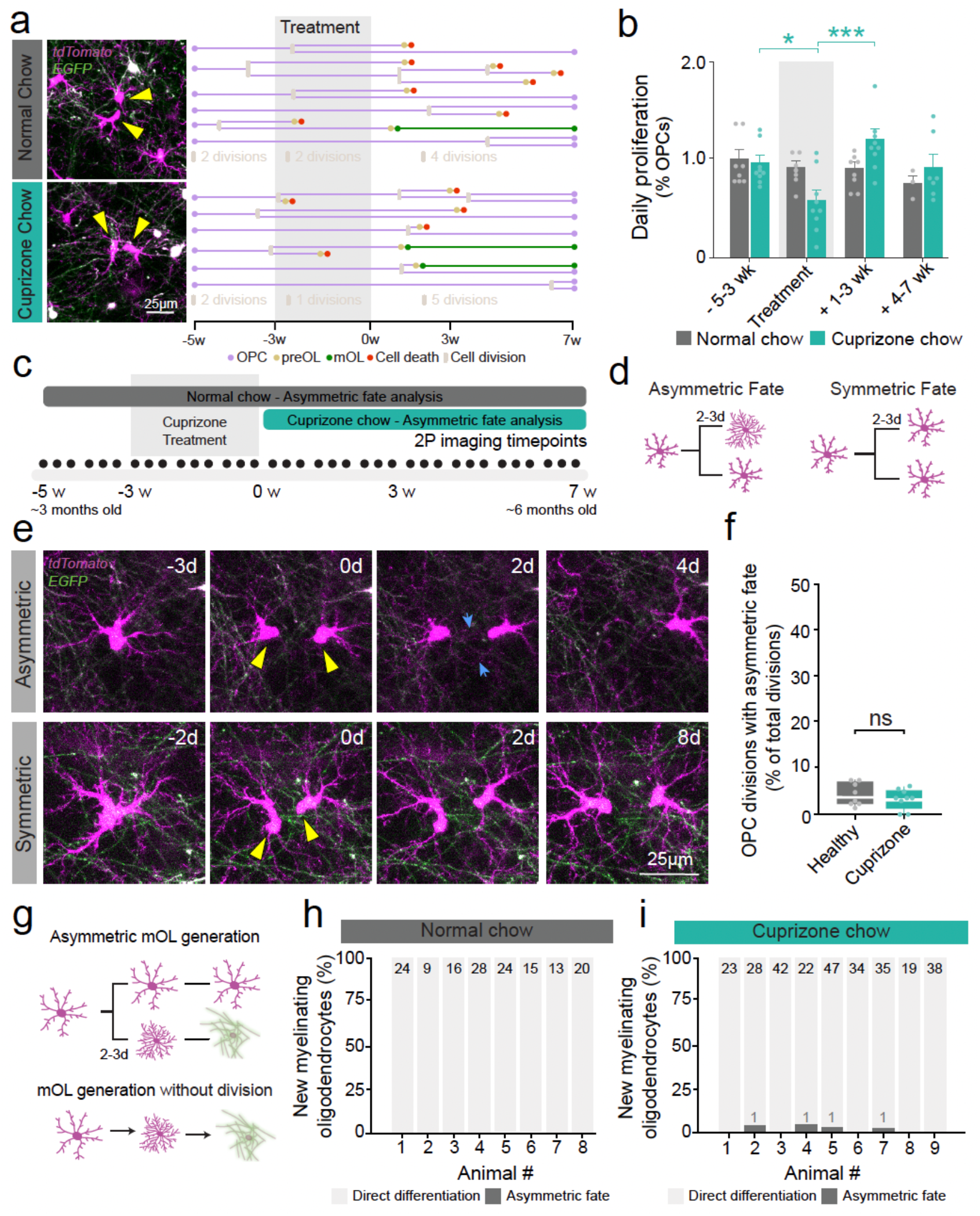
Asymmetric division does not govern the production of myelinating oligodendrocytes during regeneration. **a**, Left: Representative images of OPCs undergoing cell division in mice fed a normal diet (top) or a 0.2% cuprizone diet (bottom). Yellow arrows= cell division. Right: Representative quantitative lineage tracing diagrams from 6 individual OPCs in healthy mice (top) and cuprizone fed mice (bottom). OPC differentiation attempt (yellow), proliferation (grey), cell death (red), and new mOL generation (green). **b**, The daily rate of OPC proliferation in healthy and cuprizone fed mice (n_healthy_ = 8, n_cuprizone_ = 9). F(3, 40.07) = 6.18, p = 0.0015. Post-hoc comparisons with Tukey’s HSD. Error bars represent S.E.M. **c**, Experimental design to asses asymmetric OPC divisions in healthy mice and cuprizone fed mice. **d**, Cartoon depiction of OPC divisions resulting in asymmetric (left) or symmetric (right) fates of daughter cells. **e**, Example images showing asymmetric (top) and symmetric (bottom) fates of OPCs within two days of cell division. Yellow arrowheads= proliferation. Blue arrows= OPC territory invasion. **f**, Quantification of divisions resulting in asymmetric fate. Wilcoxon Two Sample test: |z| = 0.43, p = 0.6648. Plots denote IQR and median. **g**, Cartoon representation of an asymmetric division (top) versus mOL generation without a recent preceding division (bottom). **h, i**, Percentage of new mOL produced via asymmetric division (dark grey) or without a preceding division (light grey). Healthy mice (n=8): 0/149 new mOL generated by asymmetric division (**h**). Cuprizone mice (n = 9): 4/288 new mOL generated by asymmetric division (**i**). Dots denote individual animal means. For all analyses: n.s. not significant, *p < 0.05, **p < 0.01, ***p < 0.001. For detailed statistical information see Supplementary Table 1.

A remaining possibility is that there may be an increase in the frequency of asymmetric divisions, resulting in increased oligodendrocyte production. Similar to previous studies^46^, we considered an OPC division to be asymmetric if the fate of the daughter cells differed within 2-3 days after cell division. Using these guidelines, we assessed the frequency of divisions that resulted in asymmetric fate in healthy mice over twelve weeks and during regeneration in age-matched mice treated with cuprizone (weeks one to seven after treatment, **Figure 3C and 3D**). We found that cuprizone-mediated demyelination had no effect on asymmetric division, with 3.31 ± 0.71 % of OPC divisions having asymmetric fate in healthy mice and 4.33 ± 0.89 % of OPCs divisions having asymmetric fate in cuprizone mice (*P* = 0.6648, |z| = 0.433, Wilcoxon Two-Sample test, **Figure 3E and 3F**). Furthermore, the production of a new oligodendrocyte from asymmetric division was rare with 0/149 new oligodendrocytes produced via asymmetric division over twelve weeks in healthy mice, and 4/288 new oligodendrocytes produced via asymmetric division in seven weeks following cuprizone treatment (**Figure 3G-3I**). Overall, these data indicate that neither OPC proliferation nor asymmetric division substantially contribute to the production of new oligodendrocytes after demyelination in the adult motor cortex.

### Premyelinating oligodendrocyte survival increases to drive oligodendrocyte production after demyelination

There is robust replacement of lost oligodendrocytes after demyelinating injury. A recent study postulated that the rate of OPC differentiation remains unchanged after demyelination^27^ suggesting that another cellular mechanism, such preOL survival, underlies the increased rate of oligodendrocyte generation after demyelination. To examine if an increase in the amount of OPC differentiation attempts was responsible for the increased generation of myelinating oligodendrocytes, we assessed the number of OPCs that attempted differentiation in healthy and demyelinated mice. In line with previous results, we found that cuprizone-induced demyelination had no effect on the initiation of OPC differentiation (*P* = 0.1628, F(3,39.74)=1.8001, REML mixed model, **Figure 4B**; **S3B**). To further dissect this finding, we quantified the density of preOLs (BCAS1^high^, *Mobp*-EGFP-) via histology four days after removal of the cuprizone diet and found no difference between cuprizone mice and healthy age-matched controls (Healthy: 11.6 ± 1.33. 4d post-cuprizone: 13.8 ± 2.29, *P* = 0.4298, t(8) = 0.83, t-test, **Figure S4A and S4B**). These data indicate that increased OPC differentiation attempts were not responsible for the burst in oligodendrocyte generation during repair.

**Figure 4.**
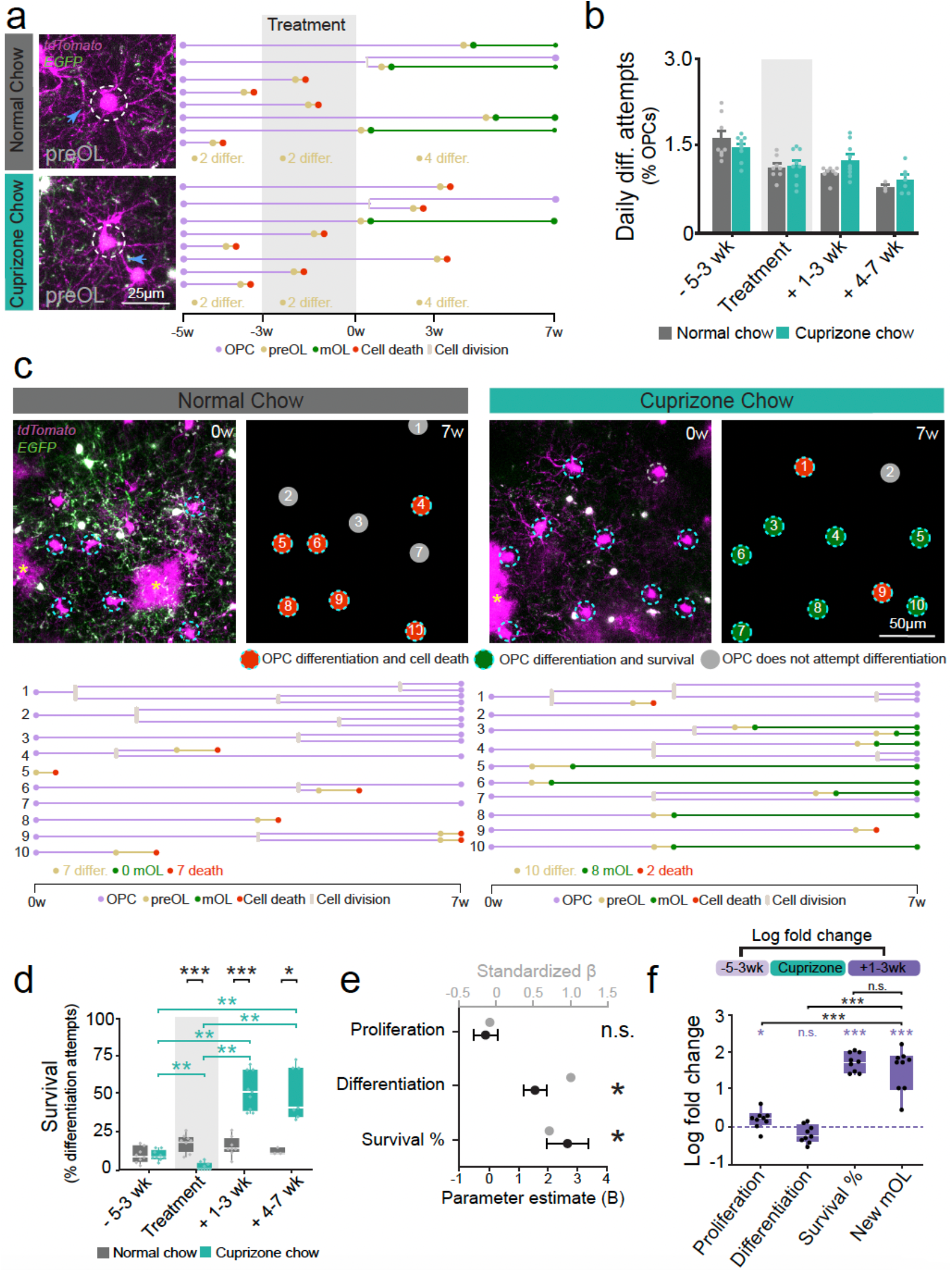
Premyelinating oligodendrocyte survival drives regeneration of myelinating oligodendrocytes. **a**, Left: Representative images showing OPC differentiation in mice fed a normal diet (top) or a 0.2% cuprizone diet (bottom). Blue arrows= OPC territory invasion. Right: Representative quantitative lineage tracing diagram from individual OPCs from healthy (top) and cuprizone fed mice (bottom). Grey box= treatment timing. OPCs attempt differentiation (yellow), proliferation (grey), cell death (red), and new mOL generation (green). **b**, Quantification of the daily rate of OPC differentiation attempts. F(3, 39.74) = 1.80, p = 0.1628. **c**, Representative images from *Olig2-CreER;R26-lsl-tdTomato;Mobp-EGFP* mice fed a normal diet (left) or a 0.2% cuprizone diet (right). Cyan dashed circles = tracked OPCs used to generate lineage tracing diagram. Colored dots (right image) = OPC fates: differentiation attempt/cell death (red), differentiation/survival (green), and no differentiation attempt (grey). Representative quantitative lineage tracing diagrams of same cells; OPC differentiation initiation (yellow), proliferation (grey), cell death (red), and new mOL generation (green). Yellow asterisk= tdTomato+ astrocytes (excluded from analysis). *Continued on next page*. **d**, Quantification of preOL survival in cuprizone fed mice vs healthy controls. Wilcoxon Two Sample test. Within group comparisons with Kruskal-Wallis test: Healthy(H(3) = 7.63, p = 0.0543), Cuprizone(H(3) = 28.10, p < 0.0001), post-hoc comparisons using Steel-Dwass method. **e**, Multiple linear regression testing the relationship between rate of mOL generation and OPC proliferation, differentiation attempts, and preOL survival. Plot denotes the Unstandardized Coefficient (B) and the Standardized Coefficient (Beta). n_mice_ = 9. Whole Model: R^2^ = 0.85, R^2^ Adj = 0.76, RMSE = 0.23, F(3,8) = 9.65, p = 0.0160. Initiation of differentiation: B = 1.55±0.40, t = 3.91, Std Beta = 1.02, p=0.0113. Proliferation: B = - 0.14±0.42, t = -0.33, Std Beta = -0.08, p = 0.7581. Survival %: B = 2.66±0.70, t = 3.81, Std Beta = 0.73, p = 0.0125. **f**, Log fold-change in the rate of cell behaviors from pre-treatment to post-treatment in mice fed cuprizone. One tailed t-test to determine if log fold-change was greater than zero: Initiation of OPC differentiation (t(8)=-2.38, p=0.9777), OPC proliferation (t(8)=2.60, p=0.0158), preOL survival (t(8)=20.78, p<0.0001), and new mOL (t(8)=8.26, p<0.0001). There is a significant effect of cell behavior on log fold-change, Kruskal-Wallis test: H(3) = 28.33, p < 0.0001. Wilcoxon rank sum with Bonferroni correction for post-hoc tests. n_mice_ = 9. Box plots represent median and IQR. Dots denote individual animal means. For all analyses: n.s. not significant, *p < 0.05, **p < 0.01, ***p < 0.001. For detailed statistical information see Supplementary Table 1.

Next, we sought to determine how the loss of myelin and oligodendrocytes affects the survival of preOLs (**Figure 4C**). During cuprizone administration, we found there is reduced survival of preOLs compared to healthy mice (Healthy: 7.18 ± 2.06 vs. Cuprizone: 1.73 ± 0.78, *P* = 0.0005, |z| = 3.45, Wilcoxon Two-Sample test; **Figure 4D**) in agreement with recent studies^16^. In contrast, during the first three weeks after removal of the cuprizone diet there was a 3.5-fold increase in the percentage of preOLs that survive the differentiation process compared to healthy mice (Healthy: 14.76 ± 2.27 % vs. Cuprizone: 51.63 ± 4.24 %, *P* = 0.0006, |z| = 3.41, Wilcoxon Two-Sample test; **Figure 4D**; **S3D**). The enhanced survival of preOLs persisted through weeks four to seven after cuprizone diet removal (Healthy: 11.58 ± 1.35 % vs. Cuprizone: 50.31 ± 6.54 %, *P* = 0.0222, |z| = 2.28, Wilcoxon Two-Sample test; **Figure 4D**). Furthermore, we found that the survival of preOLs varied across time in mice fed the cuprizone diet, whereas it remained consistent in healthy mice (Cuprizone: *P* < 0.0001. Healthy: *P* = 0.0543. Kruskal-Wallis test; **Figure 4D**; **S3D**). To further interrogate this finding, we assessed the density of BCAS1^high^, *Mobp*-EGFP-preOLs that displayed fragmented morphology (indicative of cell death) in tissue from healthy mice and tissue collected four days after removal of the cuprizone diet (**Figure S4C**). We found that there was a reduced density of preOLs with fragmented morphology four days after removal of cuprizone compared to healthy age-matched controls (Healthy: 2.6 ± 0.24 vs. 4d post-cuprizone: 0.6 ± 0.24, *P* = 0.0079, Wilcoxon Exact test; **Figure S4D**). Together, these data indicate that there is an increase in the survival of preOLs during periods of elevated oligodendrogenesis after demyelination.

To understand whether OPC proliferation, differentiation, and/or preOL survival can predict oligodendrocyte generation in young adult mice, we performed a multiple linear regression. Collectively, these three predictors account for 76.4% of the variance in oligodendrocyte generation during regeneration after cuprizone treatment (F(3,8) = 9.65, P = 0.0160). Assessing each cellular behavior’s individual, unique contributions, proliferation did not positively predict oligodendrocyte generation (B = -0.14 ± 0.42, Std Beta = -0.08, P = 0.7581), whereas preOL survival (B = 2.66 ± 0.70, Std Beta = 0.73, P = 0.0125) and OPC differentiation attempts (B = 1.55 ± 0.40, Std Beta = 1.02, P = 0.0113) both positively predict oligodendrocyte generation (**Figure 4E**). These results indicate that OPC differentiation and preOL survival together predict oligodendrocyte generation. Next, we sought to determine the extent each cellular behavior changed after cuprizone treatment. Here, we analyzed the log fold change in OPC proliferation, OPC differentiation, and preOL survival from pre-treatment to post-treatment in cuprizone treated animals. We found that there was an increase in OPC proliferation, preOL survival (Survival %), and oligodendrocyte generation (New mOL) compared to baseline, but we did not observe an increase in OPC differentiation attempts (One-tailed t-test: New mOL, *P* < 0.0001; Survival %, *P* < 0.0001; Proliferation, *P* = 0.0158; Differentiation, *P* = 0.9777; **Figure 4F**). Finally, we found that the change in preOL survival was comparable to the change in new oligodendrocyte generation, whereas the change in OPC differentiation attempts and OPC proliferation were markedly lower than the change in new oligodendrocyte generation (New mOL vs. Differentiation, |z| = 3.53, p = 0.0012; New mOL vs. Proliferation, |z| = 3.44, p = 0.0018; New mOL vs. Survival %, |z| = 0.44, p = 0.65; Wilcoxon rank sum with Bonferroni correction for post-hoc tests; **Figure 4F**). Thus, preOL survival increases after demyelination to drive the production of new oligodendrocytes.

### Impaired premyelinating oligodendrocyte survival contributes to age-related decline in oligodendrocyte production

Previous evidence suggests that preOL survival may be variable across the lifespan as survival rates differ between development and adulthood^7,9,10^. Whether preOL survival in the aging brain contributes to the age-related decline in the production of new myelinating oligodendrocytes is unclear. To assess how oligodendrocyte production, OPC differentiation, OPC proliferation, and preOL survival change throughout adulthood, we used quantitative lineage tracking in middle-aged (11-14 month old) *Olig2-CreER;R26-lsl-tdTomato;Mobp-EGFP* mice by long-term *in vivo* two-photon imaging of motor cortex every two to three days for up to 12 weeks (**Figure 5A and 5B**). In addition, we delivered 5-ethynyl-2’-deoxyuridine (EdU) via drinking water to *Mobp*-EGFP mice for 21 days and collected tissue for histological analysis after a five week chase period to assess the same cellular behaviors in young adult (6 month old), middle-aged (14 month old), and old (21 month old) mice via histology (**Figure S5A**).

**Figure 5.**
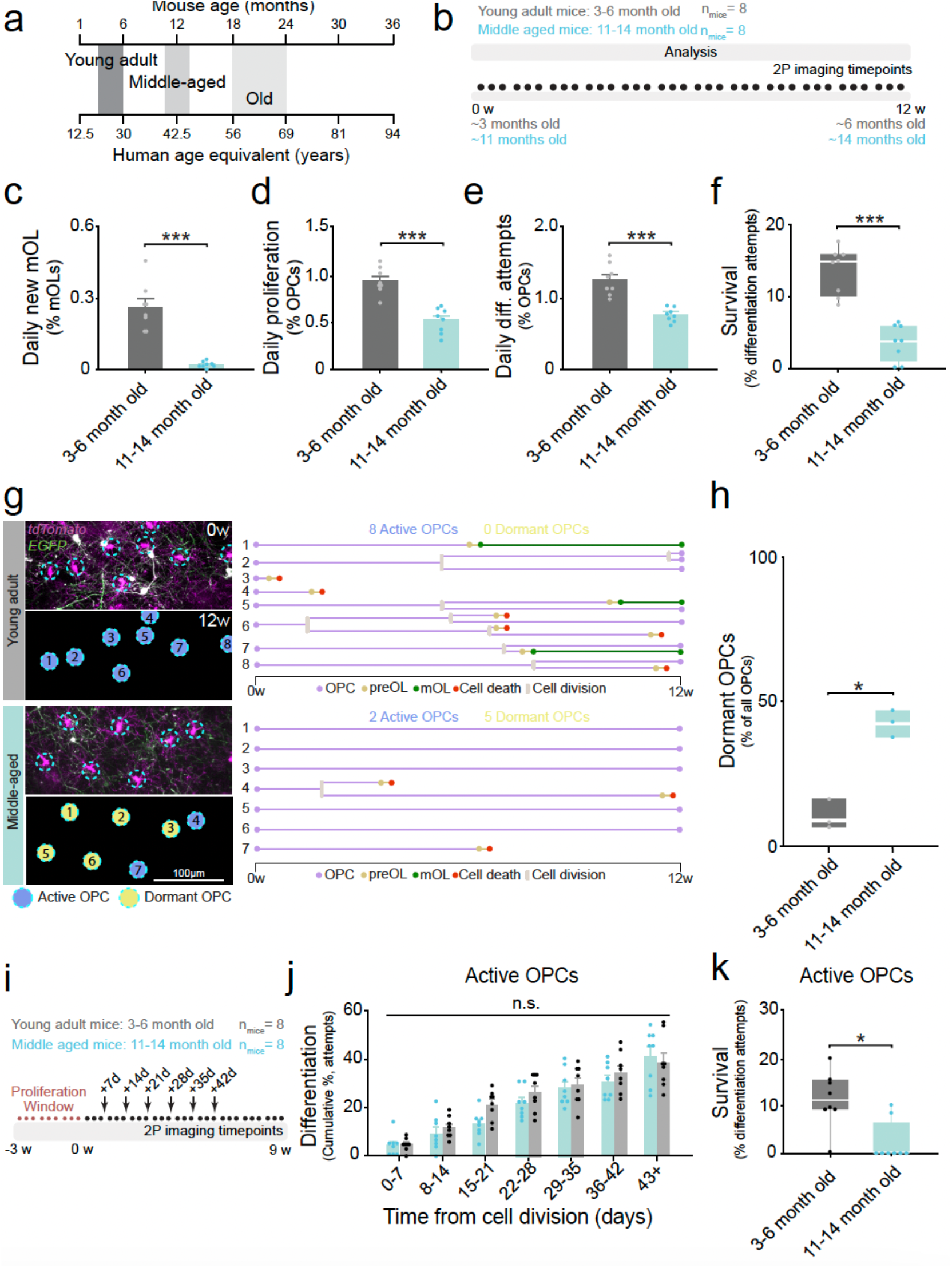
Decreased premyelinating oligodendrocyte survival contributes to age-related decline in the production of new myelinating oligodendrocytes. **a**, Schematic of mouse age relative to human age modified from^83^. **b**, Experimental design for *in vivo* imaging throughout adulthood (n_young-adult_ = 8, n_middle-aged_ = 8). Data was collected simultaneously for young adult and middle-aged cohorts; therefore, analysis was performed on the same healthy young adult mice used in previous figures. **c**, Quantification of daily rate of mOL in middle-aged versus young adult mice. Welch’s t-test: t(7.27) = -6.98, p = 0.0002. Error bars represent S.E.M. Dots denote animal means. **d**, Quantification of the daily rate of OPC proliferation in middle-aged versus young adult mice. Pooled t-test: t(14) = -6.29, p < 0.0001. Error bars represent S.E.M. **e**, Quantification of the daily rate at of OPCs differentiation attempts in middle-aged versus young adult mice. Welch’s t-test: t(8.15) = -6.50, p = 0.0002. Error bars represent S.E.M. **f**, Quantification of the preOL survival in middle-aged versus young adult mice. Wilcoxon Two Sample test: |z| = 3.31, p = 0.0009. Plots denote IQR and median. *Continued on next page*. **g**, Representative images (left) and quantitative lineage tracing diagrams (right) for OPCs from young adult (top) and middle-aged (bottom) *Olig2-CreER;R26-lsl-tdTomato;Mobp-EGFP* mice fed a normal diet. Cyan dashed circles= OPCs tracked for 12 weeks. Yellow circles= dormant OPCs; blue circles= active OPCs. Lineage diagrams: OPC differentiation initiation (yellow), proliferation (grey), cell death (red), and new mOL generation (green). **h**, Percentage of “dormant” OPCs in middle-aged vs young adult mice. Wilcoxon Exact test, p = 0.0495. n_young adult_ = 3, n_middle-aged_ = 3. Plots denote IQR and median. **i**, Experimental design to monitor OPC fate after proliferation. **j**, Quantification of cumulative percentage of OPCs differentiation attempts by week after proliferation in young adult versus middle-aged mice. Kolmogorov-Smirnov test: D(112) = 0.089, p = 0.3338. n_young adult_ = 8, n_middle-aged_ = 8. Error bars represent S.E.M. **k**, Percentage of differentiation events that result in survival for cells that underwent division during the “proliferation window” in young adult and middle-aged mice. Wilcoxon Rank Sum test: |z| = 2.57, p = 0.010. n_young adult_ = 8, n_middle-aged_ = 8. Plots denote IQR and median. Dots represent individual animal means. For all analyses: n.s. not significant, *p < 0.05, **p < 0.01, ***p < 0.001. For detailed statistical information see Supplementary Table 1.

To determine the extent of age-related decline in oligodendrogenesis, we compared the daily rate of oligodendrocyte generation in middle-aged mice to healthy young adult mice. We found that new oligodendrocyte production decreased in middle-aged mice by 92% via *in vivo* imaging (*P* = 0.0002, t(7.27) = 6.98, Welch’s t-test; **Figure 5C**). In agreement, we found that the number of new EdU+,*Mobp*-EGFP+ oligodendrocytes was reduced as mice aged via histology (*P* = 0.0002, H(2) = 17.50, Kruskal-Wallis test; **Figure S5B** and **S5C**). Specifically, we found oligodendrocyte generation was reduced by 82% in middle-aged compared to young adult mice (*P* = 0.0059, Steel-Dwass method, **Figure S5C**) and a further reduction in new oligodendrocytes in old mice (21m vs. 14m: *P* = 0.0196, 21m vs. 6m: *P* = 0.0028; **Figure S5C**).

Next, we sought to understand if a reduction in OPC proliferation or differentiation attempts contributed to the age-dependent decline in new oligodendrocyte generation. Through *in vivo* imaging, We found a 45% reduction in OPC proliferation as measured by *in vivo* imaging (*P* < 0.0001, t(14) = -6.29, t-test; **Figure 5D**). This result was also supported via histological analysis, where we found a 61% reduction in the density of PDGFRα+EdU+ OPCs in middle-aged compared to young adult mice. (*P* = 0.0002, H(2) = 17.25, Kruskal-Wallis test; **Figure S5D** and **S5E**). We found that the rate of OPC differentiation attempts was reduced by 42% in middle-aged mice compared to young adult mice (*P* = 0.0002, t(8.15) = 6.50, Welch’s t-test; **Figure 5E**). Furthermore, we found that the density of BCAS1^high^, *Mobp*-EGFP-preOLs was reduced by 43% in middle-aged compared to young adult mice via histology (*P* = 0.0012, H(2) = 13.39, Kruskal-Wallis test; **Figure S5F and S5G**). These data indicate that both OPC proliferation and differentiation are reduced as mice age.

Our *in vivo* imaging analysis revealed a 92% reduction in oligodendrocyte generation from young adulthood to middle-age; however, we only observed a 42% reduction in OPC differentiation attempts. The discrepancy between these values suggested that reduced preOL survival may also contribute to the age-dependent reduction in oligodendrocyte generation. To examine this possibility, we quantified the proportion of preOLs that survived the differentiation process in young adult and middle-aged mice. *In vivo* imaging revealed that preOL survival was reduced in middle-aged mice compared to the young adult mice (Young adult: 13.44 ± 1.15% vs. Middle-aged: 3.46 ± 0.92%, *P* = 0.0009, |z| = 3.31, Wilcoxon Two-Sample test; **Figure 5F**). In support of reduced preOL survival in aged mice, histological approaches showed an effect of age on the percentage of BCAS1^high^, *Mobp*-EGFP-preOLs that exhibited a fragmented morphology (*P* = 0.0025, H(2) = 11.95, Kruskal-Wallis test; **Figure S5H** and **S5I**). Together, these data indicate that an age-related decline in preOL survival contributes to the reduced production of new oligodendrocytes in the aging brain. Nevertheless, the concurrent reduction in OPC differentiation attempts suggests that reduced oligodendrogenesis in aged animals is due to a combination of reduced differentiation of OPCs and reduced survival of preOLs.

Previous studies show that as animals age there is an accumulation of senescence markers in the OPC population^47,48^ concurrent with reduced differentiation of OPCs into new oligodendrocytes^32,49^. We hypothesized that an increase in dormant OPCs with aging may affect the frequency of OPC differentiation and proliferation events. Thus, we sought to assess the prevalence of dormant OPCs in young adult and middle-aged mice. Similar to recent studies of neural stem cells^50^, we defined OPCs to be dormant if they did not attempt differentiation or undergo cell division over the course of 84 days (12 weeks) of longitudinal *in vivo* imaging (**Figure 5G**). To quantify the prevalence of dormant OPCs, we first determined the cellular age of individual OPCs as time from last cellular division over 12 weeks of longitudinal *in vivo* imaging (n_young adult_ = 3, n_middle-aged_ = 3, **Figure S5J**). In young adult and middle-aged mice, we found that there was a large distribution of OPC post-division age, where a majority of cells were between zero and 84 days from last division (**Figure S5K**). A small percentage of the OPC population (11%) in young adult animals did not undergo cell division or attempt differentiation over the 84 days of observation and were defined as dormant (**Figure 5H**). In contrast, we found an increase in the prevalence of dormant OPCs in middle-aged compared to young adult mice(Young adult: 10.52 ± 3.03 % vs. Middle-aged: 42.46 ± 2.88 %, *P* = 0.0495, Wilcoxon Exact test; **Figure 5H**). These data indicate that fewer OPCs in the middle-aged cortex participate in differentiation and proliferation, thereby contributing to reduced oligodendrocyte production.

Finally, we sought to determine if there were age-dependent changes in OPC differentiation attempts and preOL survival within the non-dormant, or active, OPC population. To do this, we performed a similar analysis to previous studies^46^, where we quantified the temporal dynamics of differentiation attempts for OPCs that underwent cell division within the first 21 days of *in vivo* imaging (**Figure 5I**). Within the active population, we found that the temporal dynamics of differentiation attempts after cell division did not differ between young adult and middle-aged mice (*P* = 0.3338, D(112) = 0.089, Kolmogorov-Smirnov test; **Figure 5J**), indicating that active cells are not impaired in their capacity to attempt differentiation in middle-aged mice. To assess preOL survival of active cells in young and middle-aged mice, we compared the fate of all preOLs derived from active OPCs that attempt differentiation. We found that preOL survival is reduced in the active population of OPCs in middle-aged mice compared to young adults (Young adult: 10.94 ± 2.04% vs. Middle-aged: 2.29 ± 1.51%, *P* = 0.010, |z| = -2.57, Wilcoxon Two-Sample test; **Figure 5K**). These data indicate that the temporal dynamics of differentiation attempts after cell division are unchanged for the active population of OPCs in middle-aged mice. However, despite normal initiation of differentiation, these active OPCs in middle-aged mice exhibited reduced survival at the preOL stage during oligodendrocyte differentiation.

### Premyelinating oligodendrocyte survival drives regeneration in middle-aged mice

In patients with multiple sclerosis (MS), remyelination often fails to fully restore demyelinated lesions^51^. Typically patients are not diagnosed with MS until later in life^26,52^, and previous studies show there is diminished regeneration in aged animals^28,53,54^. Our data from young adult mice showed that preOL survival increases to drive regeneration of oligodendrocytes; however, it remains unclear if preOL survival contributes to regeneration in mice whose age more closely correlates with that of human MS patients (**Figure 6A**). To examine the cellular mechanisms underlying regeneration of lost oligodendrocytes after injury in middle-aged mice, we fed 11-14 month old mice a diet containing 0.2% cuprizone to elicit demyelination and oligodendrocyte death (**Figure 6B**). We assessed OPC proliferation, differentiation attempts, preOL survival, and oligodendrocyte generation before, during, and after demyelinating injury in the middle-aged brain through quantitative lineage tracing (cuprizone treated, n_mice_ = 6, 11-14 month old; healthy mice, n_mice_ = 8, 11-14 month old). Treatment with 0.2% cuprizone diet for three weeks resulted in the loss of pre-existing mature oligodendrocytes in middle-aged mice albeit to a lesser extent than young adult animals (**Figure 6C**; Young: 69.56 ± 4.73 % vs. Middle-aged: 31.93 ± 5.08 %, *P* < 0.0001, t-test) consistent with previous work that showed that cuprizone is less effective in aged animals^55^. Following the removal of the cuprizone diet, we found a concomitant regenerative oligodendrogenesis response (**Figure 6C**; Spearman’s ρ = 0.72, p < 0.0001) that was not observed in age matched control mice fed a normal diet (**Figure S6A**). Contrary to young adult mice, we did observe the loss of some mature myelinating oligodendrocytes in healthy middle-aged mice (**Figure S6A**), potentially reflecting stochastic oligodendrocyte death observed in aged mice in previous studies^56^. Demyelinating injury resulted in a reduction in the size of the myelinating oligodendrocyte population in cuprizone treated animals that did not fully recover to healthy or pretreatment levels across seven weeks (**Figure S6B**).

**Figure 6.**
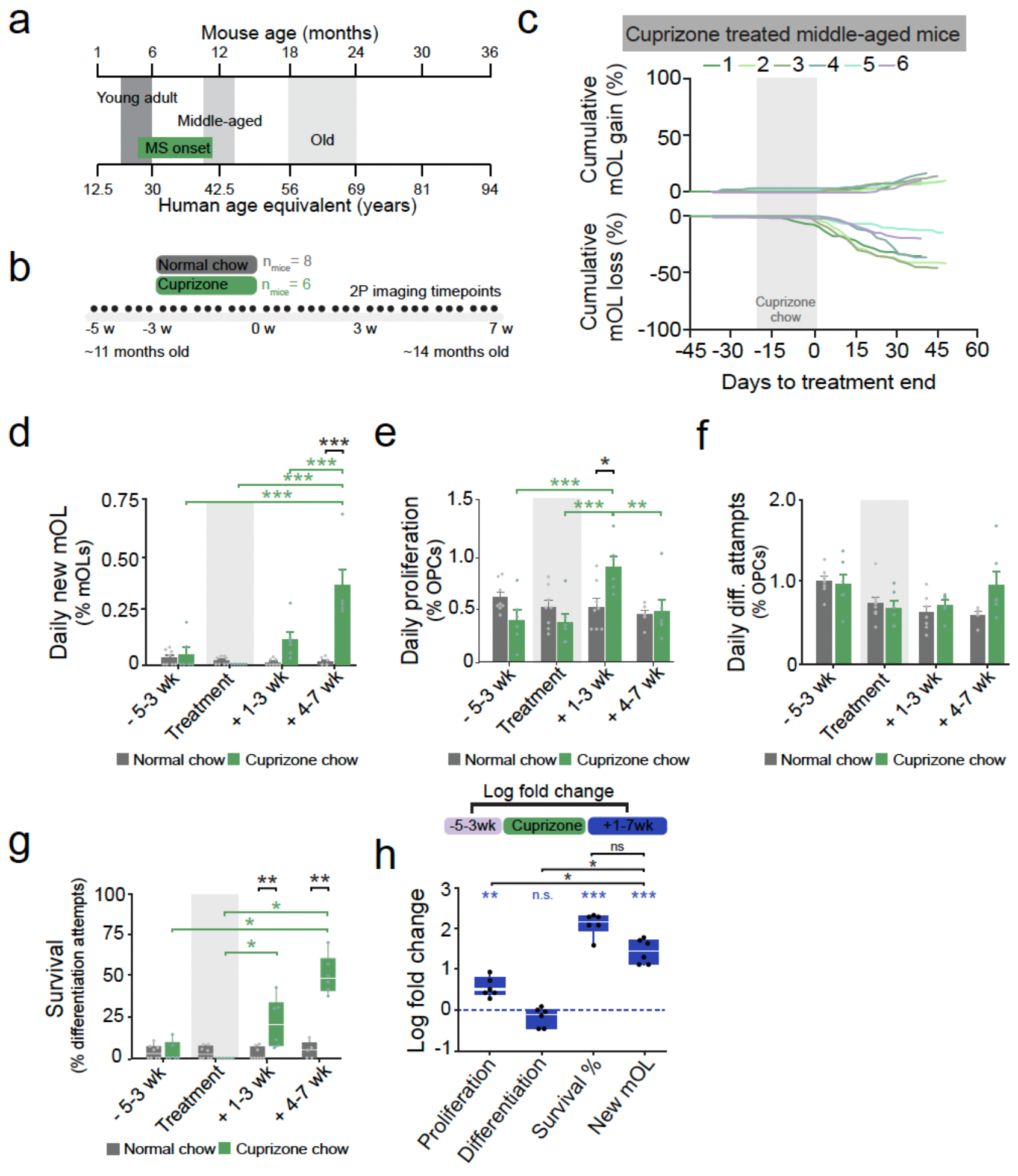
Premyelinating oligodendrocyte survival drives regeneration of oligodendrocytes in middle-aged cortex. **a**, Schematic of mouse age relative to human age modified from^83^. **b**, Experimental design *in vivo* imaging across demyelination and repair in middle-aged mice (n_healthy_ = 8, n_cuprizone_ = 6). Data was collected simultaneously for healthy and cuprizone middle-aged mice; therefore, the healthy mice are the same as in Figure 6. **c**, Cumulative oligodendrocyte gain and loss as a percentage relative to the oligodendrocyte population on day one of imaging in cuprizone mice. Color coded lines represent individual mice. Grey box= cuprizone treatment. **d**, Quantification of the daily rate of mOL generation in middle-aged mice fed cuprizone or normal chow. F(3, 35.71) = 15.70, p < 0.0001. Post hoc tests with Tukey’s HSD. n_healthy_ = 8, n_cuprizone_ = 6. Error bars represent S.E.M. **e**, Daily rate of OPC proliferation in middle-aged mice fed cuprizone or normal chow. F(3, 34.3) = 8.38, p = 0.0003. Post-hoc tests with Tukey’s HSD. n_healthy_ = 8, n_cuprizone_ = 6. Error bars represent S.E.M. **f**, Daily rate of OPC differentiation attempts in middle-aged mice fed cuprizone or normal chow. F(3, 34.01) = 2.44, p = 0.0807. n_healthy_ = 8, n_cuprizone_ = 6. Error bars represent S.E.M. **g**, Quantification of preOL survival in middle-aged mice fed cuprizone or normal chow. Wilcoxon Rank Sum test: -5-3wk (|z| = 0.07, p = 0.943), Treatment(|z| = 1.86, p = 0.0625), +1-3wk(|z| = 2.64, p = 0.0082), +4-7wk(|z| = 2.65, p = 0.0080). Kruskal-Wallis test: Healthy(H(3) = 0.48, p = 0.9213), Cuprizone(H(3) = 19.25, p = 0.0002). Post-hoc comparisons using Steel-Dwass method. n_healthy_ = 8, n_cuprizone_ = 6. Box plots represent median and IQR. **h**, Log fold-change in the rate of cell behaviors from pre-treatment to post-treatment in middle-aged mice fed cuprizone. One tailed t-test to determine if log fold-change was greater than zero: Initiation of OPC differentiation (t(5)=-1.65, p=0.9197), OPC proliferation (t(5)=5.75, p=0.0011), preOL survival (t(5)=59.74, p<0.0001), and new mOL (t(5)=11.77, p<0.0001). Kruskal-Wallis test: H(3) = 17.33, p = 0.0006. Wilcoxon rank sum with Bonferroni correction for post-hoc tests. n_mice_ = 6. Box plots represent median and IQR. Dots denote individual animal means. For all analyses: n.s. not significant, *p < 0.05, **p < 0.01, ***p < 0.001. For detailed statistical information see Supplementary Table 1.

To assess the dynamics of oligodendrocyte production across demyelination and repair, we divided the experiment into four phases (**Figure 2**). We found that generation of oligodendrocytes was modulated by cuprizone treatment (*P* < 0.0001, F(3, 35.71) = 15.70, REML mixed model; **Figure 6D**). Specifically, oligodendrocyte generation was increased during weeks four to seven following cuprizone compared to healthy age matched controls (*P* < 0.0001, Tukey’s HSD test; **Figure 6D**). Moreover, the daily rate of oligodendrocyte generation within the group of demyelinated mice was increased during weeks four to seven after removal of the cuprizone treatment compared to all other phases of the experiment (**Figure 6D**). The increase in oligodendrocyte generation was delayed compared to young adult mice (**Figure 2E**), possibly due to the comparative delay in the loss of myelinating oligodendrocytes in middle-aged mice. These data indicate oligodendrocyte regeneration is preserved in the middle-aged brain.

In our analysis of healthy middle-aged mice, we found an increased prevalence of dormant OPCs relative to young adult mice (**Figure 5H**) and it is possible that dormant OPCs are recruited to participate in regeneration in the middle-aged cortex. To examine this possibility, we compared the prevalence of dormant OPCs after 12 weeks of imaging in middle-aged mice treated with cuprizone compared to healthy age-matched controls. We found that there was no difference in the percentage of OPCs that exhibited a dormant phenotype between healthy or demyelinated middle-aged mice (Middle-aged healthy: 42.46 ± 2.88% vs. Middle-aged cuprizone: 36.22 ± 3.25%, *P* = 0.2286, Wilcoxon Exact test; **Figure S6C**). Thus, demyelination in middle-aged mice fails to recruit dormant OPCs to participate in regeneration.

Next, we sought to determine whether more OPCs proliferate or attempt differentiation during regeneration compared to healthy conditions in middle-aged mice. We found that OPC proliferation was increased during weeks one to three after treatment compared to healthy age-matched controls and within the group of mice treated with cuprizone (*P* = 0.0459, Tukey’s HSD test; **Figure 6E**). In contrast and in alignment with data from young adult animals (**Figure 4B**), there was no effect of cuprizone treatment on OPC differentiation attempts within or across groups for the entire experimental time course (*P* = 0.0807, F(3, 34.01) = 2.44, REML mixed model; **Figure 6F**). These data indicate that OPCs in the middle-aged brain do not respond to demyelinating injury by increasing differentiation attempts, but they do respond with an increased rate of division in the weeks immediately following removal of cuprizone treatment.

To understand how the loss of myelin and oligodendrocytes affects the survival of preOLs in the middle-aged brain, we analyzed the fate of all preOLs generated from OPCs that attempted differentiation across demyelination and repair. We found that survival of preOLs varied across time in middle-aged mice fed the cuprizone diet, but not within healthy control mice (Cuprizone: *P* < 0.0002. Healthy: *P* = 0.9213. Kruskal-Wallis test; **Figure 6G**). Specifically, we found an increase in preOL survival beginning during the first three weeks after removal of the cuprizone diet (Healthy: 2.62 ± 1.32% vs. Cuprizone: 21.44 ± 6.18%, *P* = 0.0082, Wilcoxon Two-Sample test), an effect that persisted into weeks four to seven post-cuprizone during which we observed a 10-fold increase in preOL survival compared to healthy age-matched controls (Healthy: 4.70 ± 2.32% vs. Cuprizone: 50.41 ± 4.80%, *P* = 0.0080, Wilcoxon Two-Sample test; **Figure 6G**).

To further dissect this finding and determine which cellular behavior most influenced the regeneration of oligodendrocytes in middle-aged mice, we analyzed the log fold change in OPC proliferation, differentiation, and preOL survival from pre-treatment to post-treatment in cuprizone animals and compared them to the log fold change of new oligodendrocyte generation. We found that there was an increase in OPC proliferation, preOL survival (Survival %) and oligodendrocyte generation (New mOL) compared to baseline, but we did not observe an increase in OPC differentiation attempts (One-tailed t-test: New mOL, *P* < 0.0001; Survival %, *P* < 0.0001; Proliferation, *P* = 0.0011; Differentiation, *P* = 0.9197; **Figure 6H**). Finally, we found the change in preOL survival was comparable to the change in new oligodendrocyte generation, whereas the change in OPC differentiation attempts and proliferation were markedly lower compared to the change in new oligodendrocyte generation (New mOL vs. Differentiation, *P* = 0.0366, |z| = 2.50; New mOL vs. Proliferation, *P* = 0.0366, |z| = 2.50; New mOL vs. Survival %, *P* = 0.1101, |z| = 2.08; **Figure 6H**). Thus, preOL survival increases after demyelination to drive the production of new oligodendrocytes in middle-aged mice.

### GPR17 antagonism enhances myelin repair via increased preOL survival

Previous work has identified a number of compounds that enhance OPC differentiation, however, these candidates have had limited success in clinical trials^38^. However, our data indicated that regeneration following a demyelinating injury is driven by preOL survival rather than increased OPC differentiation (**Figure 4**). Therefore, we sought to identify a compound that could enhance preOL survival and integration. Oligodendrocyte lineage specific G-coupled protein receptor 17 (GPR17) has been postulated as a key negative regulator of preOL survival, as GPR17 knockout mice show enhanced remyelination after demyelinating injury^41,57,58^. Furthermore, we found that GPR17 protein is expressed in BCAS1-positive preOLs (**Figure 7A**). Thus, we aimed to test the effect of Myro-02, a novel, selective, and brain penetrant GPR17 antagonist, on oligodendrocyte regeneration and preOL survival in the context of demyelinating injury. We elicited demyelination and oligodendrocyte death by treating three month old *Olig2-CreER;R26-lsl-tdTomato;Mobp-EGFP* mice a 0.2% cuprizone diet for three weeks, followed by three weeks of twice-daily oral gavage of vehicle or 12mg/kg Myro-02 (24mg/kg/day) (**Figure 7B**). We monitored oligodendrocyte loss and regeneration as well as reconstructed cell lineages of individual OPC clones via longitudinal two-photon *in vivo* imaging acquired three times per week before, during, and after cuprizone administration in mice treated with vehicle (n_mice_= 8) or Myro-02 (n_mice_ = 8) (**Figure 7B**).

**Figure 7.**
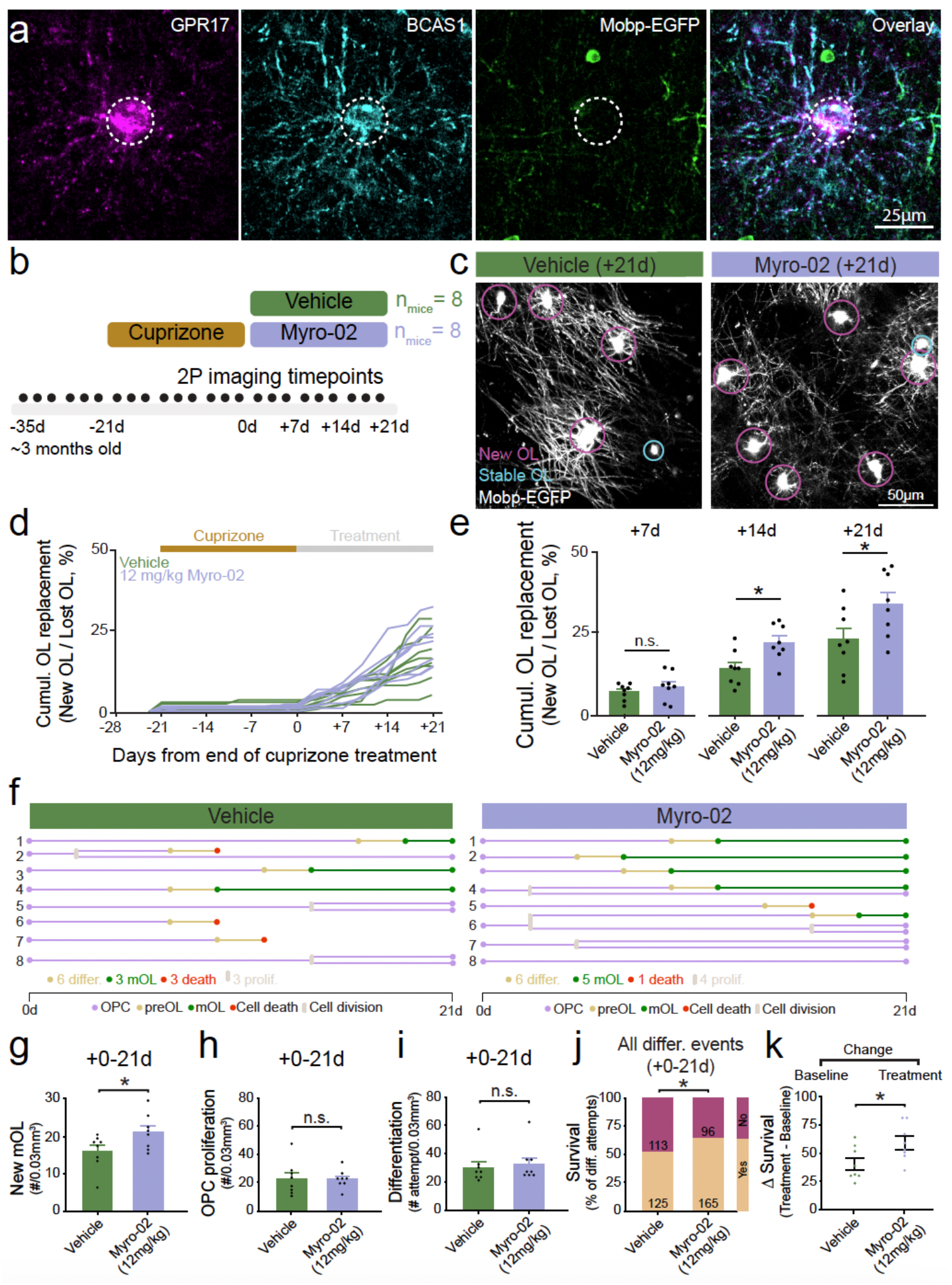
GPR17 antagonist enhances regeneration via increased premyelinating oligodendrocyte survival. **a**, Representative image of BCAS1^high^ *Mobp*-EGFP-preOL co-labeled with GPR17 from a 4 month old *Mobp*-EGFP mouse. **b**, Experimental design for *in vivo* imaging across demyelination and repair in 3-6 month old mice treated with Vehicle or Myro-02 (n_vehicle_ = 8, n_Myro-02_ = 8). **c**, Representative images of new mOL generation on day 21 of treatment in mice treated with Vehicle or Myro-02. Magenta circles= mOL generated during treatment. Cyan circles= stable mOLs generated prior to treatment. **d**, Cumulative oligodendrocyte replacement as a percentage. Grey box= treatment timing. Lines represent individual mice. **e**, Cross sectional measurements of cumulative oligodendrocyte replacement percent. Day 7: Pooled t-test, t(14) = 0.94, p = 0.3646. Day 14: Pooled t-test, t(14) = 2.96, p = 0.0104. Day 21: Pooled t-test, t(14) = 2.20, p = 0.0453. **f**, Representative quantitative lineage tracing diagrams of OPC clones tracked over the course of 21 days of treatment in Vehicle (left) or Myro-02 (right) treated mice. OPC differentiation attempts (yellow), proliferation (grey), cell death (red), new mOL generation (green). **g**, Quantification of new mOL generation during treatment with Myro-02 or Vehicle. Pooled t-test: t(14) = 2.15, p = 0.0498. **h**, Quantification of OPC proliferation during treatment with Myro-02 or Vehicle. Pooled t-test: t(14) = 0.03, p = 0.9798. **i**, Quantification of OPC differentiation attempts during treatment with Myro-02 or Vehicle Pooled t-test: t(14) = 0.47, p = 0.6457. **j**, Percentage of preOL survival during treatment with Myro-02 or Vehicle. Myro-02: 165/261 preOLs survived differentiation, Vehicle: 125/238 preOLs survived differentiation. p = 0.0156, 𝒳^2^=5.85, Pearson Chi-Square. **k**, Change in preOL survival during treatment with Myro-02 or Vehicle relative to baseline. Pooled t-test: t(14) = 2.38, p = 0.0319. Error bars denote S.E.M. Dots denote individual animal means. For all analyses: n.s. not significant, *p < 0.05, **p < 0.01, ***p < 0.001. For detailed statistical information see Supplementary Table 1.

Consistent with age-matched untreated mice (**Figure 2C**), oligodendrocyte loss continued three weeks following removal of cuprizone treatment in both Myro-02 and vehicle treated mice (**Figure S7A**). Oligodendrocyte loss was consistent across groups and over the three week treatment period (+7d: Vehicle: 31.53 ± 5.44% vs. Myro-02: 33.74 ± 6.12%; +14d: Vehicle: 53.98 ± 7.02% vs. Myro-02: 51.42 ± 6.07%; +21d: Vehicle: 64.36 ± 6.12% vs. Myro-02: 61.96 ± 5.82%, **Figure S7B**). Upon cuprizone diet removal, we found a characteristic burst of new oligodendrocyte regeneration (**Figure S7C)**. As the generation of oligodendrocytes is correlated with the extent of oligodendrocyte loss (*P* = 0.0417, R^2^=0.526, and **Figure S7F**), we assessed oligodendrocyte replacement, calculated as the number of oligodendrocytes generated during the three week treatment period as a proportion of total oligodendrocyte loss (**Figure 7D and 7E**)^14^. We found that Myro-02 enhanced cumulative oligodendrocyte replacement relative to vehicle at 14 days (Vehicle: 14.15 ± 1.869% vs. Myro-02: 22.23 ± 1.97%, *P* = 0.0104, t-test, **Figure 7E**) and 21 days (Vehicle: 23.32 ± 2.44% vs. Myro-02: 34.14 ± 3.52%, *P* = 0.0453, t-test, **Figure 7E**) after the onset of treatment. These results indicate that GPR17 antagonism, via Myro-02 treatment, increased oligodendrocyte regeneration following cuprizone-mediated demyelination.

To investigate the cellular principles underlying the enhanced repair induced by GPR17 antagonism, we assessed the total number of oligodendrocytes generated, and OPCs that proliferated or attempted differentiation during treatment (0-21days following end of cuprizone treatment) (**Figure 7F**). In accordance with enhanced oligodendrocyte replacement, we found that the total number of oligodendrocytes generated during treatment was increased by Myro-02 compared to vehicle treatment (Vehicle: 15.63 ± 1.62 vs. Myro-02: 20.63 ± 1.68, *P* = 0.0498, t-test, **Figure 7G**) whereas we found no differences in total OPC proliferation (Vehicle: 22.13 ± 4.25 vs. Myro-02: 22.00 ± 2.31, **Figure 7H**) or OPC differentiation attempts (Vehicle: 29.75 ± 4.14 vs. Myro-02: 32.63 ± 4.50, **Figure 7I**). Thus, neither changes in OPC differentiation nor proliferation account for the observed increase in oligodendrocyte generation.

Similar to age-matched untreated mice (**Figure 4F**), both vehicle and Myro-02 treated mice increased preOL survival to a level that was comparable to the increase in oligodendrocyte generation after cuprizone treatment, indicating that preOL survival, not OPC differentiation attempts, led to the generation of new oligodendrocytes (**Figure S7D and S7E**). To continue to explore if there was a relationship between Myro-02 treatment and preOL survival, we conducted a contingency analysis of OPC differentiation attempts and found preOLs from mice treated with Myro-02 were more likely to survive than preOLs from mice treated with vehicle, suggesting that Myro-02 is associated with increased survival of preOLs (*P* = 0.0156, 𝒳^2^=5.85, Pearson Chi-Square, **Figure 7J**). Due to high inter-mouse variability, next we assessed the change in percentage of preOL survival during the treatment period normalized to baseline on a mouse-by-mouse basis. We found that Myro-02 increased preOL survival to a greater extent than vehicle treatment (Vehicle: 40.03 ± 5.24 % vs. Myro-02: 58.59 ± 5.76 %, P = 0.0319, t-test, **Figure 7K**). In support of this result, using this same analytical approach we found that while Myro-02 increased the change in generation of oligodendrocytes to a greater extent than vehicle treatment, there was no difference in the change of OPC differentiation attempts or proliferation between Myro-02 and vehicle treated mice (**Figure S7G-I**). Altogether, these data indicate that Myro-02 enhances repair following a demyelinating injury by promoting preOL survival to increase oligodendrocyte regeneration.

## Discussion

Our findings identify premyelinating oligodendrocyte (preOL) survival—not the initiation of OPC differentiation—as the principal cellular regulator of adult oligodendrogenesis. Although OPCs continuously attempt differentiation in the healthy brain^27^, the generation of mature oligodendrocytes is remarkably inefficient^10^. Using quantitative lineage analyses to simultaneously track OPC proliferation, differentiation, and fate over time, we demonstrate that preOL survival is dynamically tuned across physiological states and ages (summarized in **Figure S8**). In young adult cortex, preOL survival is low, consistent with a tight regulation of new oligodendrocyte formation. Following demyelination, preOL survival rises dramatically, enabling rapid restoration of oligodendrocyte number despite the unchanged frequency of precursor differentiation attempts. Conversely, in the aging brain, we find that preOL survival is reduced, even within the population of non-dormant OPCs, contributing to age-dependent reduction in oligodendrogenesis. Finally, we show that GPR17 antagonism promotes the survival of preOLs, which in turn increases the replacement of lost oligodendrocytes after demyelination. These results demonstrate preOL survival dynamically changes in response to demyelination and that preOL survival can be therapeutically targeted to enhance regeneration.

These results support a working model in which preOL survival functions as a form of “gain control” on oligodendrocyte output. Rather than solely relying on upregulation of OPC differentiation initiation, which is relatively inflexible and time consuming, the adjustment of the fraction of preOLs that stably integrate into circuits allows for rapid, fine-tuned alterations to new myelin formation. This architecture provides a means to match oligodendrocyte production to environmental demand, enabling sharp increases in output during regeneration and tight restriction during homeostasis. Such gain control may also intersect with experience-dependent myelination. While previous work showed that neural activity and motor learning increased OPC proliferation^59,60^ and differentiation attempts^14,61^, during development, sensory experience modulated the survival and integration of preOLs^5,62^. Trophic signaling via growth factors promote oligodendrocyte survival^63^ and recent work shows that BDNF-TrkB signaling in OPCs is necessary for activity dependent myelination^64^, raising the possibility that signaling through TrkB receptors may modulate survival during the differentiation process. Together, these findings suggest that multiple forms of extrinsic signals may converge on preOL survival as a checkpoint to adjust myelin remodeling.

Previous studies show that OPCs form functional synapses with neurons^65,66^, and that these synaptic connections are maintained throughout the process of cell division^67^. These bona-fide synaptic connections are lost early during the differentiation from OPC to preOL ^68,69^; however, there are emerging roles for synaptic proteins in the development and stabilization of nascent myelin sheaths^70,71^. Our data show that preOL survival is modulated by demyelination, suggesting that newly available axons, and their synaptic connections, may influence preOL survival. Furthermore, we observed that most new oligodendrocytes are produced from the direct differentiation of OPCs that have not recently divided, not from asymmetric division. These data indicate that the survival of preOLs may rely on the establishment of precise and targeted relationships with axons within an individual cell’s territory, and that reestablishing these connections after cell division is essential for the stable integration of preOLs into neuronal circuitry. Regional comparisons further support this idea: white matter and the developing CNS, both of which provide rich axonal and trophic environments, exhibit higher rates of oligodendrogenesis than adult gray matter ^31,44,72^. Other work shows that preOL survival is higher during development than in adulthood^7,9,10^. These lines of evidence suggest that the cue(s) controlling survival of preOLs may be found by investigating differences between the developing and adult brain or by comparing white and gray matter in the adult brain.

At the molecular level, we identify GPR17 signaling as an effective regulator of the preOL survival checkpoint. GPR17 is maximally expressed at the preOL stage, negatively regulates differentiation, and can regulate oligodendrocyte maturation^5,58,73^. Studies have also postulated GPR17 as a key negative regulator of preOL survival through inducing expression of the pro-apoptotic gene Xaf1^57^. We found that antagonizing GPR17 with Myro-02 enhances preOL survival *in vivo* without altering differentiation attempts, demonstrating that survival can be pharmacologically modulated independently of other lineage transitions. GPR17 ligands, including cysteinyl leukotrienes, increase after injury and inflammation^74,75^, suggesting this pathway may represent a context-dependent brake on regeneration. Whether GPR17 antagonism will promote broader effects on oligodendroglia lineage transitions in more restrictive or highly inflammatory injury models^40,41^ will require additional study. However, injury alone increases preOL survival even without GPR17 inhibition, implying that additional pro-survival programs are engaged during repair. Knockout of transcription factor EB (TFEB) causes ectopic myelination via enhanced survival of preOLs during differentiation^12^, leading to the possibility that extrinsic modulation of this signaling pathway could influence survival of preOLs during differentiation. Alternatively, known modulators of oligodendrocyte production, such as GPR37 signaling ^76^ or the retinoid receptor RXR-γ^35^, could promote survival during differentiation. Other potential pathways include extracellular matrix signaling, cell-cell communication with astrocytes and microglia, as well as interactions with neuronal axons^77^. Future studies should address whether these cues promote the initiation of OPC differentiation, preOL survival, or both.

Modulation of preOL survival may have therapeutic implications for diseases with myelin damage, for example multiple sclerosis and other neurodegenerative disease with myelin deficits such as Alzheimer’s disease. Although remyelination failure in MS is often attributed to impaired OPC recruitment or differentiation^78^, the presence of OPCs and preOLs within some lesions suggests that survival barriers may also limit regeneration^17,24,26,79^. Inflammatory lesion environments may suppress preOL survival, rendering strategies that solely promote the initiation of OPC differentiation insufficient. Combining pro-differentiation and pro-survival interventions may therefore be necessary to restore effective oligodendrocyte replacement.

Aging presents the related but distinct challenge of OPC senescence, with impaired initiation of OPC differentiation, and myelin repair deficits in aged animals^47,52,53^. While middle-aged OPCs include a growing population of dormant, likely senescent cells, non-dormant OPCs maintain normal differentiation kinetics, yet still produce fewer oligodendrocytes due to poor preOL survival. These findings suggest that both the aged microenvironment and intrinsic OPC biology^48^ impose dominant barriers to myelin repair. Encouragingly, preOL survival in non-dormant OPCs can be enhanced after demyelination, indicating that preOL survival remains responsive to injury-associated signals. Previous studies have found that the age-related decline in differentiation potential can be rescued in part by treatment with Metformin^32^. However, it is unclear if compounds such as Metformin or Myro-02 modulate preOL survival in the middle-aged brain or alternatively affect OPC senescence. Future work should define how aging alters local trophic support, axonal signaling, inflammation, and extracellular matrix composition, and whether interventions that modify these features, such as senolytics, metabolic modulators, or anti-inflammatory strategies, can enhance preOL survival *in vivo*. Together, our results reframe adult oligodendrogenesis around a central, regulatable survival checkpoint of the premyelinating oligodendrocyte. These findings may open new avenues for modulating myelin plasticity and regeneration across health, disease, and aging.

## Materials and Methods

*Animals*. All animal experiments were conducted in accordance with protocols approved by the Animal Care and Use Committee at the University of Colorado Anschutz Medical Campus. Male and female mice used in these experiments were kept on a 14-h light–10-h dark schedule with ad libitum access to food and water. All mice were randomly assigned to conditions and were age-matched across experimental groups. Generation and genotyping of BAC transgenic lines *Mobp-EGFP*^80^ (Gensat), *Olig2-CreER*^81^ and tdTomato reporter mouse line^82^ (Ai14, Allen Brain Institute) have been previously described. For experiments using the inducible Cre–loxP system to label oligodendrocyte lineage cells or dividing cells, mice were injected with tamoxifen (see “Tamoxifen injection” section below for details).

### Two-photon microscopy

Cranial windows were prepared as previously described^6,10,14^. Briefly, mice (2-3 month old or 10-11 month old) were anesthetized with isoflurane (induction, 5%; maintenance, 1.5–2.0%, mixed with 0.5 L/min O2), and their body temperature was maintained at 37 °C with a thermostat-controlled heating plate. The skin over the right cerebral hemisphere was removed and the skull cleaned. A 2 × 2 mm region of skull over the motor cortex (0–2 mm anterior to bregma and 0.5–2.5 mm lateral) was removed using a high-speed dental drill. For cranial windows, a piece of cover glass (VWR, No. 1) was placed in the craniotomy and sealed with dental cement (C&B Metabond) and a custom metal plate with a central hole was attached to the skull for head stabilization. *In vivo* imaging sessions began after a minimum of 3 weeks (chronic cranial window). During imaging sessions, mice were anesthetized with isoflurane and immobilized by attaching the head plate to a custom stage. Images were collected using a Zeiss LSM 7MP microscope equipped with a BiG GaAsP detector using a mode-locked Ti:sapphire laser (Coherent Ultra) tuned to 920 nm. The average power at the sample during imaging was < 30 mW. Vascular landmarks were used to identify the same cortical area over longitudinal imaging sessions. Image stacks were 425 μm ×425 μm × 336 μm (1,024 ×1,024 pixels; corresponding to layers I–III) from the cortical surface.

### Cuprizone-mediated demyelination

Cortical demyelination was induced as described previously^14^. Briefly, demyelination was induced in 3 month old or 11 month old *Olig2-CreER;R26-lsl-tdTomato;Mobp-EGFP* mice using 0.2% cuprizone (Sigma Chemical, C9012), stored in a glass desiccator at 4°C. Cuprizone was mixed into powdered chow (Harlan) and fed to mice in custom feeders for 3 weeks on an ad libitum basis. Feeders were refilled every 2–3 days, and fresh cuprizone chow was prepared weekly. Cages were changed weekly to avoid build-up of cuprizone chow in bedding, and to minimize reuptake of cuprizone chow following cessation of diet via coprophagia. Mice were returned to normal diet after 3 weeks of feeding the 0.2% cuprizone diet.

### Preparation and delivery of Myro-02

Myro-02 solution was prepared at a concentration of 1.2mg/mL in a vehicle solution of 0.5% w/v 400cP Methylcellulose + 2% v/v Tween-80 in dH_2_O. Mice were delivered Myro-02 or vehicle twice daily via oral gavage at a dose of 12 mg/kg (10 mL/kg solution). To prepare the vehicle solution, 2 g of 400cP Methylcellulose (MC, Spectrum Chemical ME136, 9004-67-5) was put into 400 mL of dH_2_O and stirred overnight at room temperature. The following day, 8 mL of MC/dH_2_O solution was discarded and replaced with 8 mL Tween-80 (T-80, MP Bio 02103170-CF). Vehicle solution was stored at 4 °C and used throughout the study. The compound was formulated into a homogenous suspension, by adding Myro-02 to the appropriate volume of vehicle in a glass vial to achieve a final concentration of 1.2 mg/mL. Compound was stored at room temperature for up to 7 days. 10 minutes before dosing, compound was stirred to ensure homogeneity. Scientists were blinded to the identity of compound and vehicle for the duration of dosing.

### Immunohistochemistry

Mice were anesthetized with an intraperitoneal injection of sodium pentobarbital (100 mg per kg body weight) and transcardially perfused with 4% paraformaldehyde in 0.1 M phosphate buffer (pH 7.0–7.4). Brains were postfixed in 4% paraformaldehyde for 2-4 h at 4 °C, transferred to 30% sucrose solution in PBS (pH 7.4) and stored at 4 °C for at least 24 h. Brains were extracted, frozen in TissuePlus OCT and sectioned coronally at 30-μm thick. Immunostaining was performed on free-floating sections. Next, if necessary, sections underwent antigen retrieval where they were incubated for 2 minutes at 95°C in preheated 10mM Tris, 1mM EDTA, and 0.05% Tween 20 in PBS, pH 9. After sections had cooled to room temperature, they were incubated in blocking solution (5% normal donkey serum, 0.3% Triton X-100 in PBS, pH 7.4) for 1–4 h at room temperature, then incubated overnight at 4°C in primary antibody (listed along with secondary antibodies in **Supplementary Table 2**). Secondary antibody incubation was performed at room temperature for 2 h. Sections were mounted on slides with Vectashield antifade reagent (Vector Laboratories). Images were acquired using a laser-scanning confocal microscope (Nikon A1R). Image analysis was done using the Cell Counter plugin on Fiji. Briefly, a minimum of 3 FOV’s were analyzed for each animal for each measurement, each FOV was 614.88µmx614.88µm spanning L1-L5 of the primary motor cortex.

### Tyramide signal amplification

Tyramide signal amplification (TSA) was performed for all GPR17 immunostaining. Briefly, sections were incubated in 1% hydrogen peroxide solution for 40 minutes at room temperature and washed 3 times for 5 minutes in PBS. Sections were then incubated in blocking solution (1% bovine serum albumin (BSA), 0.3% Triton X-100 in PBS, pH 7.4) for 30 minutes at room temperature before incubating in GPR17 antibody (listed along with secondary antibodies in **Supplementary Table 2**) cocktail overnight at 4°C. Sections were then washed three times for 5 minutes in PBS and incubated with a peroxidase conjugated secondary antibody for 30 minutes at room temperature before being washed three times for 5 minutes in PBS. Sections were then incubated with a fluorophore tyramide (AZDye 546 Tyramide, CCT-1539-1, Vector Laboratories) diluted 1:100 in a 0.0015% hydrogen peroxide solution for 5 min. After incubation in fluorophore tyramide solution, sections were washed three times for 5 minutes in PBS and additional staining was performed as described above.

### Image processing and analysis

Image processing and analysis was done as previously described^6,10,14^. Briefly, image stacks were analyzed using Fiji or ImageJ. All analyses were performed on unprocessed images. When generating figures, image brightness and contrast levels were adjusted for clarity. For pseudocolor display of individual OPCs, a maximum intensity projection of the cell of interest was generated and manually colorized in Adobe Photoshop. Longitudinal image stacks were registered using the Fiji plugins Correct 3D drift or PoorMan3Dreg. Blinding to experimental conditions was used for analyzing image stacks from two-photon imaging.

### Cell tracking

Custom Fiji scripts were written to follow oligodendrocytes and OPCs in four dimensions by identifying tdTomato+ and/or EGFP+ cells bodies at each time point, recording *x, y* and *z* coordinates, and defining cellular behavior (new, dying, proliferating, differentiating, or stable cells). Mature oligodendrocyte and OPC migration, proliferation, death and differentiation were defined as previously described^6,10,14^. In *Olig2-CreER;R26-lsl-tdTomato;Mobp-EGFP* mice direct death of OPCs was not observed as was previously observed in other mouse lines^6,10,14^, death oligodendroglia was only observed after presentation of stereotyped morphological changes that occur after initiation of terminal differentiation. For cell division, the timepoint that two daughter cells could first be distinguished was marked as the time of division. For initiation of differentiation, the timepoint of cell morphological change (invasion of neighboring cells or EGFP expression or cell disappearance) was marked as the time of differentiation, for ∼90% of cells this was the timepoint preceding disappearance or EGFP expression. For new myelinating oligodendrocyte formation, the timepoint of first EGFP expression was marked as the time of oligodendrocyte generation. For myelinating oligodendrocyte death, the timepoint of last observed EGFP expression was marked as the time of cell death. The area of clarity of longitudinally imaged cranial windows varies from animal to animal, and cellular tracking was limited only to the portion of the window that remained clear for the duration of imaging. Therefore, there was variation in the size of the cortical volume analyzed in each animal. To control for the variation in the size of the analyzed area, and the number of cells analyzed within that area in each animal, the daily rates of OPC and oligodendrocyte cell behaviors relative to the baseline cell number were calculated. Survival was measured as the number of cells that survived the differentiation process (co-expression of EGFP) divided by the number of OPCs that attempt differentiation during a given phase of the experiment. Equations used to calculate cell behaviors can be found below:

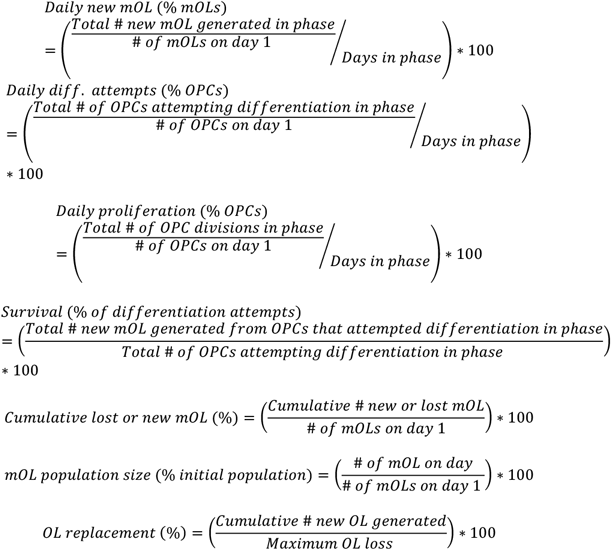

Creation of individual cell lineages. Output from the custom cell tracking Fiji script contained the time point of every OPC division, OPC differentiation (morphology), death during differentiation (cell disappearance) and survival and integration (co-expression of EGFP). Using this information, a data set was created with the cells post-division age (either day 1 of imaging, or the day of cell division), cell behavior (initiation of differentiation, cell division, disappearance, or EGFP expression), and time of cell behavior for each individual cell. With this dataset, individual cell lineages were curated manually from the tracking data.

### Cell morphology analysis

First, individual cells were identified in longitudinal image stacks based on morphological criteria described above and defined as either being an OPC or a preOL. Then the cell was cropped to contain all XYZ information using a custom Fiji macro to only include processes belonging to the cell of interest. For convex hull analysis, cells were converted to greyscale 8-bit images, max projected, and auto-thresholded using the Fiji Default algorithm. Finally, the thresholded images were analyzed using the “Hull and Circle” ImageJ plugin (https://imagej.net/ij/plugins/hull-circle.html). For Sholl analysis, cells were cropped using the same methodology above. Next, using a custom written Fiji macro the cells were converted to greyscale 8-bit format, max-projected then auto-thresholded. Sholl analysis was carried out on the thresholded cell using the ImageJ Sholl Analysis plugin. The starting radius was set to 0.5μm and the end radius set to 100μm; 0.5μm increments were used for analysis.

### Tamoxifen injections

Tamoxifen (Sigma; T5648–1G; CAS:10540– 29–1) was weighed and dispersed in sunflower seed oil (Sigma; S5007; CAS: 8001–21–6) for a final concentration of 10 mg/mL. The solution was vortexed vigorously and then sonicated for 5 min. This cycle was repeated three to four times, until tamoxifen was completely dissolved. The tamoxifen/oil emulsion was stored at 4 °C, protected from light, and used for a maximum of 7 days. Mice 4-6 weeks old were injected intraperitoneally with 100 mg/kg bodyweight of the tamoxifen/oil emulsion once per day for 5 consecutive days to induce Cre recombination.

### EdU labeling and detection

For cumulative labeling experiments EdU was delivered as previously described^31^. Briefly, EdU (Lumiprobe) was dissolved in the drinking water at 0.2 mg/ml. The water was exchanged every 48 hr and mice were exposed to EdU for 21 days followed by a 5 week chase at which point tissue was collected for analysis. EdU was visualized using a homemade mix immediately following the blocking step of the immunohistochemistry protocol. The mix was prepared by sequentially adding the following components: 909 μL PBS, 80 μL of Copper(II) sulfate pentahydrate (209198, Sigma Aldrich; 10 mM fresh stock, final concentration 800 μM), 1 μL Fluor-Azide (200 μM stock, final concentration 200 nM, Vector labs (CCT-1300-1)), and 10 μL freshly made L-Ascorbic acid (100 mg/mL frozen stock, final concentration 5.7 mM). 500 μL of the EdU mix was added to each sample, and the samples were incubated at RT for 30 minutes, covered from light. Additional staining was performed afterwards.

### Statistics

Summarized statistical information is present in the legend of each figure. Detailed descriptions of statistical tests can be found in **Supplementary Table 1**. Sample sizes were not predetermined but were comparable to relevant publications^10,14,44^. Experimental groups were replicated in multiple batches with multiple experimental groups per batch. If possible, littermates were divided into different experimental groups within the same batch. Due to the longitudinal nature of the study and the clarity of cranial windows which animals would produce full datasets could not be predicted when assigning to experimental groups. Therefore, not all experimental groups contain animals with full datasets.

All statistical analyses were performed in JMP (SAS). When data was not normality distributed, nonparametric tests were used, nonparametric tests were also used when comparing population proportions, such as “Survival (% of differentiation events)”. When data were not normally distributed and sample sizes were small (n<5 / group) the Wilcoxon Exact test was used. When comparing three or more groups, the Kruskal-Wallis test with Steel-Dwass method for post-hoc comparisons was used.

If the data were normally distributed, the variance between different groups was compared with the Brown-Forsythe test. If variances were unequal, a Welch’s t-test was performed. When variance was equal, w tests including two-tailed Pooled t-tests or analysis of variance (ANOVA) with Tukey’s honestly significant difference (HSD) post-hoc tests were performed. Two-tailed tests and α ≤ 0.05 were always employed unless otherwise specified. To investigate the relationship between two variables, simple linear regression was used; the coefficient of determination (R2) and the p-value of an ANOVA testing the null-hypothesis that there is no linear relationship between the tested variables were reported.

To test for interactions between treatment and experimental phase, statistical mixed modeling was performed, a restricted maximum likelihood approach with an unbounded variance component and least-squared Tukey’s HSD post-hoc tests was used. All models were two fixed effects with full factorial models. Batch effects were statistically controlled for. For all models, ‘Mouse ID’ was a random variable, and this random variable was nested within cohort. For data visualization, all error bars represent the standard error of the mean, all bar graphs denote means and all box plots illustrate medians and IQRs unless otherwise specified. For all analyses: n.s. not significant, *p < 0.05, **p < 0.01, ***p < 0.001.

## Acknowledgements

We thank Ryan Mettetal for machining expertise, Andrew Scallon and the CU Anschutz Optogenetics and Neural Engineering Core (P30NS048154) for 3D printing and stage design, past and current members of the Hughes Lab and the Colorado Glia (COG) gatherings for discussion. We thank D. Lecca and M. Abbracchio (Università degli Studi di Milano) for graciously sharing the antibody to GPR17. This investigation was supported by the National Institutes of Health NINDS (NS115975, NS125230, NS132859, NS134829) to EGH and in part by Fast Forward commercial research funding award (FF-2312-42544) from the National Multiple Sclerosis Society to Myrobalan Therapeutics, Inc..

## Author Contributions

M.E.S. and E.G.H. conceived the project. M.E.S. designed and performed the experiments, analyzed the data, and generated the figures. M.W. and Y.M.C. assisted with performing experiments and data analysis for **Figure 7** and **Figure S7**. A.K. assisted with data analysis for **Figure 5, Figure S5, Figure 6**, and **Figure S6**. S.K. assisted with data analysis for **Figure S1**. T.A.S. provided technical expertise and compound for experiments conducted in **Figure 7** and **Figure S7**. C.M., L.A.O., and K.H.S. performed surgeries. G.D.F.N. performed part of the in vivo imaging. A.R.C. performed confocal imaging and genotyping. E.G.H. supervised the project. M.E.S. and E.G.H. wrote the manuscript.

