## Supplemental Figures for "Premyelinating Oligodendrocyte Survival Governs CNS Remyelination"

### Supplementary Video Legends

**Supplementary Video 1 | Oligodendrocyte precursor cell division.**
*In vivo* time series in a *Olig2-CreER; R26-lsl-tdTomato;Mobp-EGFP* mouse showing a OPC (magenta) clone that undergoes 3 divisions over 89 days of imaging. Note that the two daughter cells migrate apart after division. Division is denoted by yellow arrows and occurs on day 25, day 50, and day 71. Images were acquired in layer 1 of the motor cortex. The white dashed circle denotes the cell of interested. Cell stage is written in the top left. Imaging day is denoted in the top right. Colors: tdTomato (magenta), *Mobp*-EGFP (green).

**Supplementary Video 2 | Stable oligodendrocyte precursor cell.**
*In vivo* time series in a *Olig2-CreER; R26-lsl-tdTomato;Mobp-EGFP* mouse showing a stable OPC (magenta). Note the motile processes and individual territory of the highlighted OPC. Images were acquired in layer 1 of the motor cortex. The white dashed circle denotes the cell of interested. Cell stage is written in the top left. Imaging day is denoted in the top right. Colors: tdTomato (magenta), *Mobp*-EGFP (green).

**Supplementary Video 3 | Initiation of OPC differentiation and the survival and integration of a preOL.**
*In vivo* time series in a *Olig2-CreER; R26-lsl-tdTomato;Mobp-EGFP* mouse showing an OPC (magenta) that differentiates into a preOL, which then survives and integrates as a myelinating oligodendrocyte. Images were acquired in layer 1 of the motor cortex. The white dashed circle denotes the cell of interested and blue arrows highlight territory invasion of neighboring OPCs after differentiation into a preOL. Note the development of long, slender, ramified processes at the preOL stage on day 19 and day 21. Survival and integration occurs by day 25 as denoted by expression of *Mobp*-EGFP and the formation of myelin sheaths. The new oligodendrocyte then remains stable until the end of the imaging period on day 62. Cell stage is written in the top left. Imaging day is denoted in the top right. Colors: tdTomato (magenta), *Mobp*-EGFP (green).

**Supplementary Video 4 | Initiation of OPC differentiation and cell death of a preOL.**
*In vivo* time series in a *Olig2-CreER; R26-lsl-tdTomato;Mobp-EGFP* mouse showing an OPC (magenta) that differentiates into a preOL, which then undergoes cell death. Images were acquired in layer 1 of the motor cortex. The white dashed circle denotes the cell of interested and the blue arrow highlights territory invasion of a neighboring OPC after differentiation into a preOL on day 14. Note the development of long, slender, ramified processes at the preOL stage on day 14. On day 16, there is cellular fragmentation of the cell of interest. The cell is lost, presumedly to cell death, between day 14 and day 16. Cell stage is written in the top left. Imaging day is denoted in the top right. Colors: tdTomato (magenta), *Mobp*-EGFP (green).

### Supplementary Figures

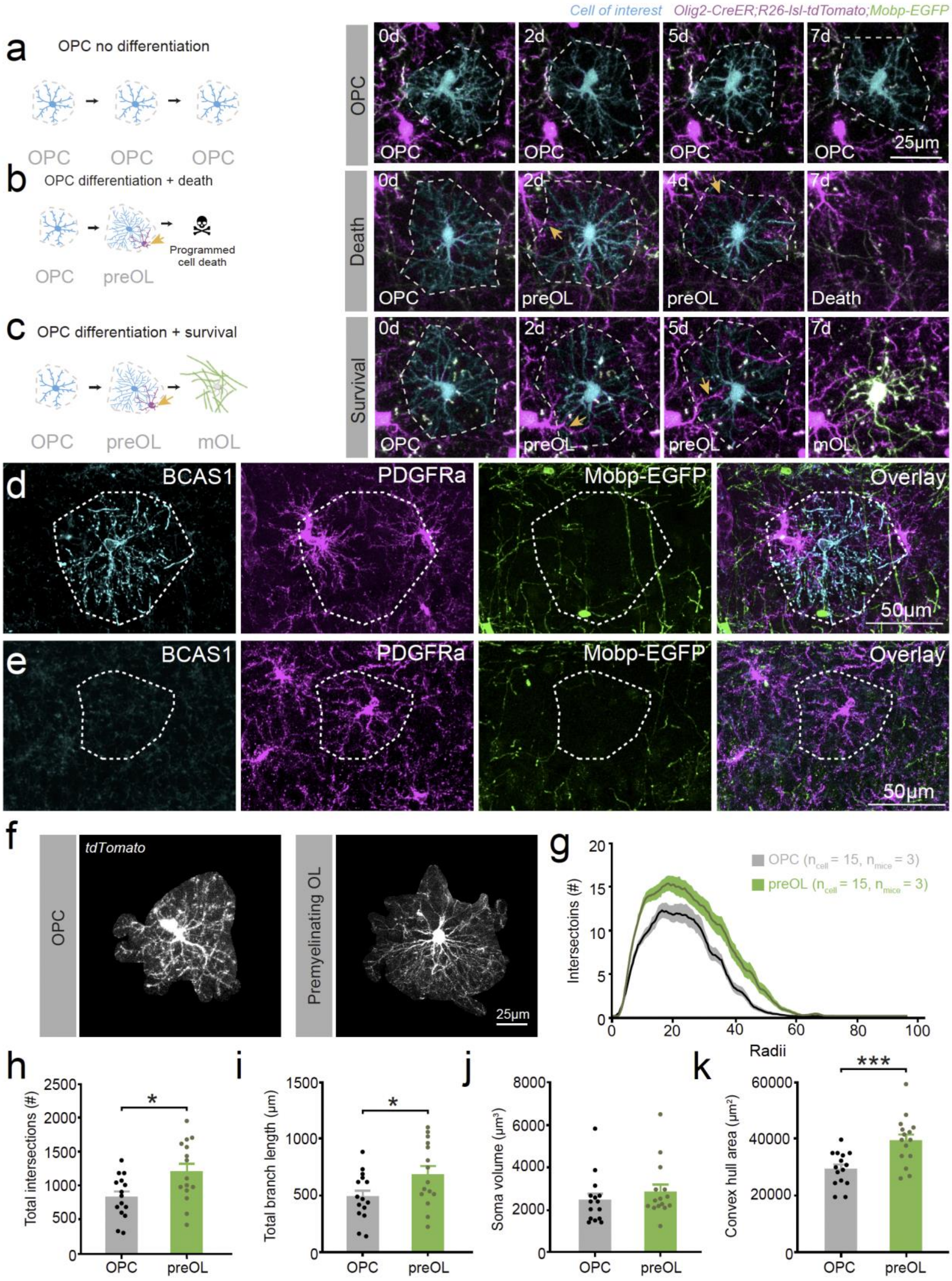


**Supplementary Figure 1**

**Supplementary Figure 1 | Identification of premyelinating oligodendrocytes in *Olig2-CreER;tdTomato;Mobp-EGFP* mice.**

**a, b, c,** Representative images of individual OPCs from longitudinal *in vivo* imaging of *Olig2-CreER;R26-lsl-tdTomato;Mobp-EGFP* mice. **a,** Territory of an OPC that does not attempt differentiation or undergo cell division. **b**, Territory of an OPC that differentiates into a preOL and then undergoes programmed cell death. **c**, Territory of an OPC that differentiates into a preOL, survives and integrates as a new mOL. **a-c:** Cell of interest= cyan, grey dashed line= cell's territory, orange arrows= neighboring OPCs territory invasion. **d,** Representative images from a *Mobp*-EGFP (green) mouse highlighting PDGFR$\alpha$+ (magenta) OPCs invading the territory of a Bcas1$^{high}$ (cyan) *Mobp*-EGFP- preOL. White dashed line= preOL's cellular territory. **e**, Representative images from a *Mobp*-EGFP (green) mouse highlighting PDGFR$\alpha$+ (magenta) OPCs exhibiting homotypic repulsion to maintain exclusive cellular territories from neighboring OPCs. White dashed line = OPC territory. **f,** Representative images from *Olig2-CreER;R26-lsl-tdTomato;Mobp-EGFP* mice showing a z-projection of an OPC and preOL that were identified and cropped from *in vivo* images. OPCs and preOLs were *Mobp*-EGFP- and identified using the characteristic morphological changes that occur during differentiation. **g,** Average Sholl analysis plot of cells identified as OPCs or preOLs from *in vivo* images (OPCs: $n_{cell}$ = 15, $n_{mice}$ = 3. PreOLs: $n_{cell}$ = 15, $n_{mice}$ = 3). Line represents mean of all cells and shaded area represents 95% confidence interval. **h,** Quantification of total intersections from a Sholl analysis of preOLs versus OPCs. Pooled t-test, $t(28) = 2.65$, $p=0.0131$. OPCs: $n_{cell}$ = 15, $n_{mice}$ = 3. PreOLs: $n_{cell}$ = 15, $n_{mice}$ = 3. **i,** Quantification of total branch length from a Sholl analysis of preOLs versus OPCs. Pooled t-test, $t(28) = 2.23$, $p=0.0337$. OPCs: $n_{cell}$ = 15, $n_{mice}$ = 3. PreOLs: $n_{cell}$ = 15, $n_{mice}$ = 3. **j,** Quantification of soma size in OPCs versus preOLs. Pooled t-test, $t(28) = 0.92$, $p=0.3633$. OPCs: $n_{cell}$ = 15, $n_{mice}$ = 3. PreOLs: $n_{cell}$ = 15, $n_{mice}$ = 3. **k**, Cell size as measured by the area of a convex hull in preOLs versus OPCs. Pooled t-test, $t(28) = 3.76$, $p=0.0008$. OPCs: $n_{cell}$ = 15, $n_{mice}$ = 3. PreOLs: $n_{cell}$ = 15, $n_{mice}$ = 3. Dots represent individual cells, plots denote mean and S.E.M. For all analyses: n.s. not significant, *$p < 0.05$, **$p < 0.01$, ***$p < 0.001$. For detailed statistical information see Supplementary Table 1.

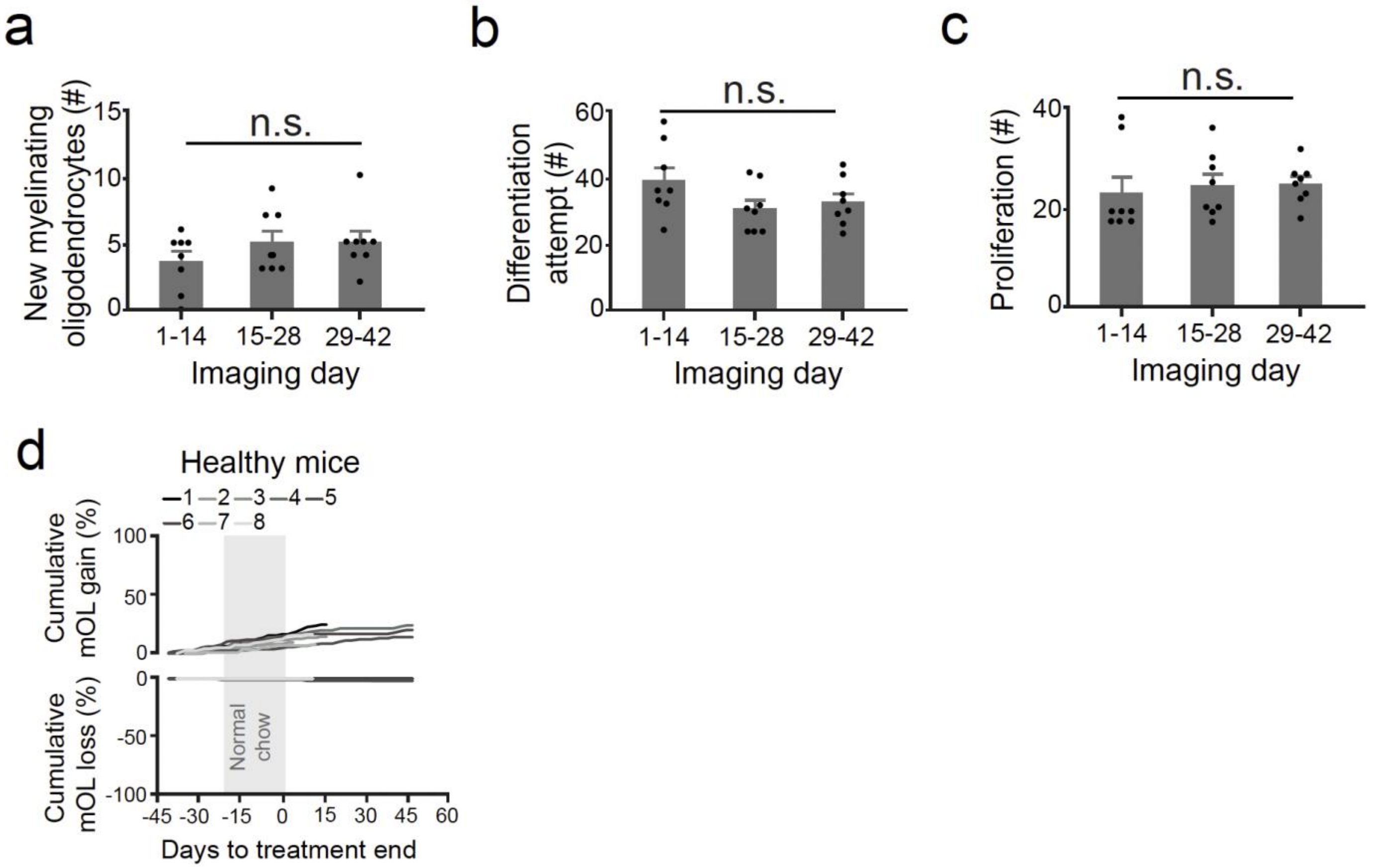


**Supplementary Figure 2**

**Supplementary Figure 2| Oligodendrocyte gain, OPC differentiation attempts, and OPC proliferation are not modulated by time**

**a,** Quantification of new mOLs generated over time in healthy young adult mice. $F(2,23) = 1.0$, $p=0.3847$, ANOVA. $n_{mice} = 8$. Dots represent animal means. **b**, Quantification of OPC differentiation attempts over time in healthy young adult mice. $F(2,23) = 1.98$, $p=0.1631$, ANOVA. $n_{mice} = 8$. **c,** Quantification of OPC proliferation over time in healthy young adult mice. $F(2,23) = 0.23$, $p=0.7939$, ANOVA. $n_{mice} = 8$. Dots represent individual animal means. Plots denote mean and S.E.M. For all analyses: n.s. not significant, *p < 0.05, **p < 0.01, ***p < 0.001. For detailed statistical information see Supplementary Table 1. **d**, Cumulative oligodendrocyte gain and loss as a percentage relative to the oligodendrocyte population on day one of imaging in healthy mice.

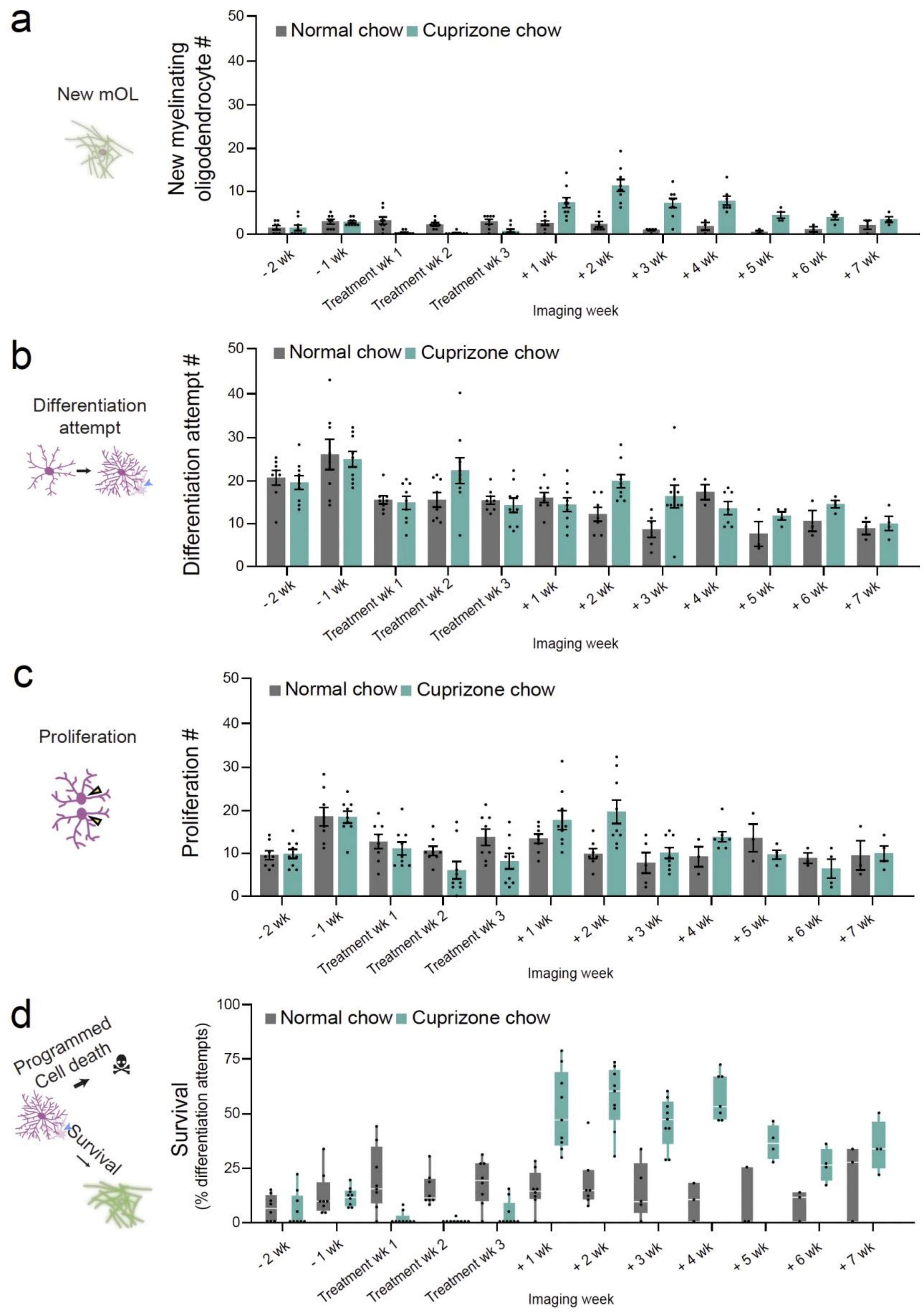


**Supplementary Figure 3**

**Supplementary Figure 3 | Raw counts of new myelinating oligodendrocytes, OPC differentiation attempts, OPC proliferation, and premyelinating oligodendrocyte survival.**

**a**, Quantification of new mOLs produced during the indicated week. Error bars, S.E.M. **b**, Quantification of OPCs that attempt differentiation during the indicated week. Error bars, S.E.M. **c,** Quantification of OPCs that underwent cell division during the indicated week. Error bars, S.E.M. **d,** Percentage of differentiation events that resulted in survival during the indicated week. Box plots represent median and IQR. Dots denote individual mice means. For all analyses: n.s. not significant, $*p < 0.05$, $**p < 0.01$, $***p < 0.001$. For detailed statistical information see Supplementary Table 1.

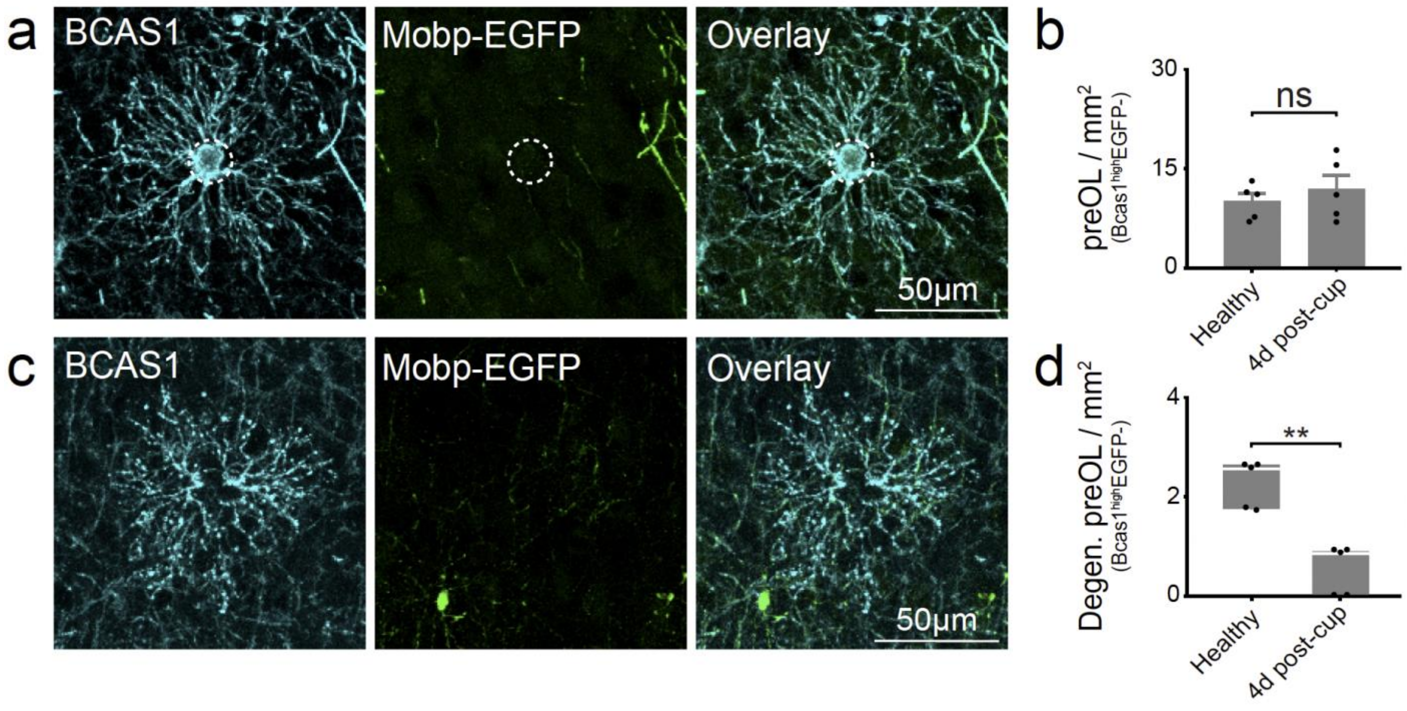


**Supplementary Figure 4**

**Supplementary Figure 4 | Oligodendroglia cell behaviors during regeneration after cuprizone treatment.** **a,** Example image from an *Mobp*-EGFP mouse showing a Bcas1$^{High}$EGFP- preOL. White dashed circle denotes cell of interest. **b**, Quantification of density of Bcas1$^{high}$*Mobp*-EGFP- preOLs in mice 4 days post-cuprizone versus healthy mice. $n_{healthy}$ = 5, $n_{cuprizone}$ = 5. Pooled t-test, t(8) = 0.83, p=0.4298. Plots denote mean and S.E.M. **c**, Example image from an *Mobp*-EGFP mouse showing a degenerating Bcas1$^{High}$ *Mobp*-EGFP- preOL. Note the fragmenting Bcas1+ processes. White dashed circle denotes cell of interest. **d**, Quantification of the density of degenerating Bcas1$^{High}$ *Mobp*-EGFP- preOLs in healthy versus 4 day post-cuprizone mice. $n_{healthy}$ = 5, $n_{cuprizone}$ = 5. Wilcoxon exact test, p=0.0079. Box plots represent median and IQR. Dots represent individual mice. For all analyses: n.s. not significant, *p < 0.05, **p < 0.01, ***p < 0.001. For detailed statistical information see Supplementary Table 1.

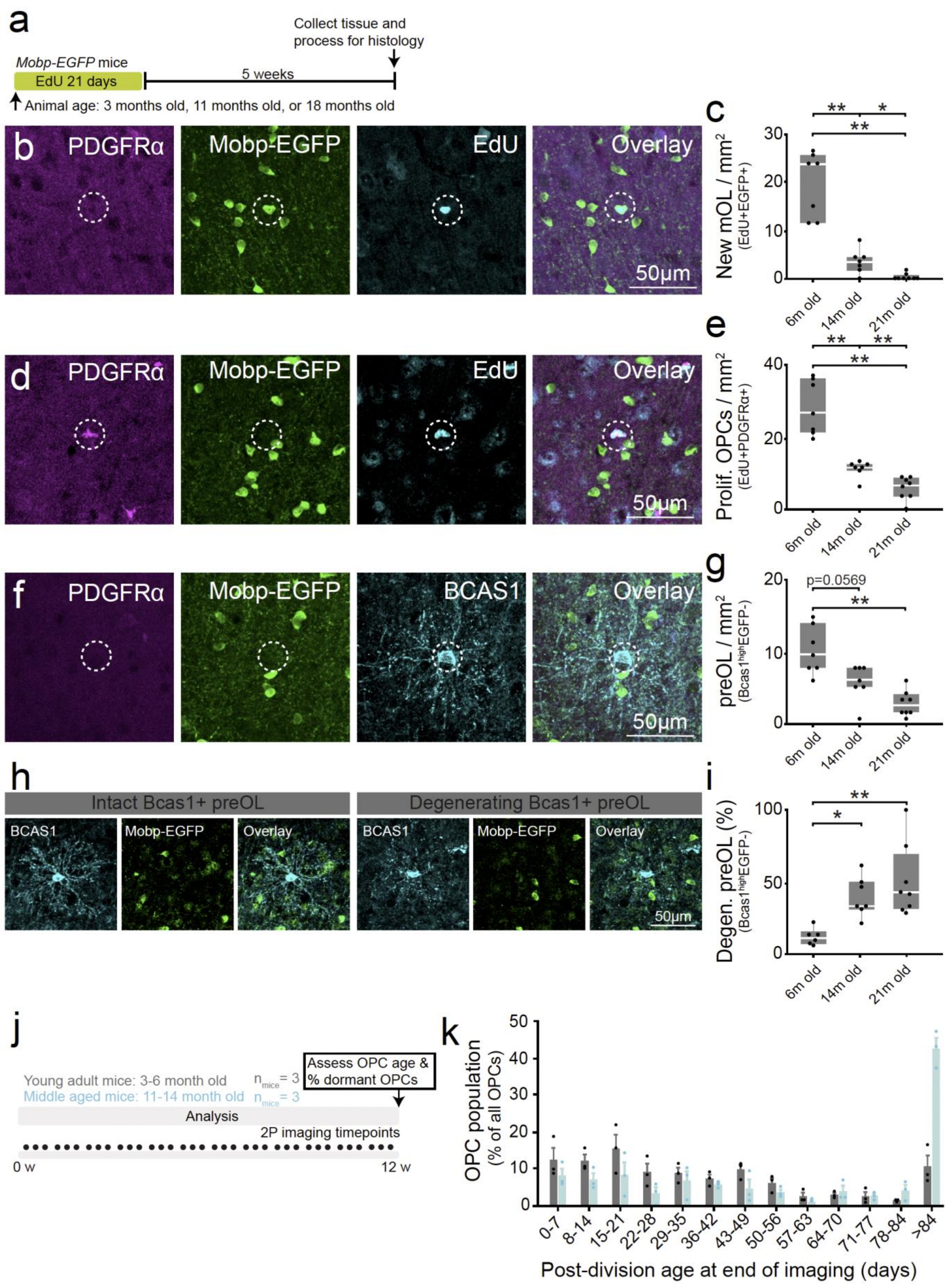


**Supplementary Figure 5**

**Supplementary Figure 5 | Cellular dynamics of oligodendroglia vary in the young adult, middle-aged, and old aged cortex.**

**a**, Experimental protocol for assessing new oligodendrocyte production, preOL density, OPC proliferation, and degenerating preOL density in the cortex of young adult (6 month old), middle-age (14 month old), and old-age (21-month -old) mice. **b,** Example image from a *Mobp*-EGFP mouse showing a newly generated myelinating oligodendrocyte (*Mobp*-EGFP+EdU+). White dashed circle denotes cell of interest. **c,** The density of *Mobp*-EGFP+EdU+ new myelinating oligodendrocytes is modulated by age. $n_{6\ month\ old}$ = 7, $n_{14\ month\ old}$ = 7, $n_{21\ month\ old}$ = 8. Kruskal-Wallis test, H(2) = 17.5, p=0.0002. Post-hoc comparisons via Steel-Dwass method. Dots represent individual mice. Box plots represent median and IQR. **d,** Example image from a *Mobp*-EGFP mouse showing a PDGFR$\alpha$+EdU+ OPC. White dashed circle denotes cell of interest. **e,** Quantification of density of PDGFR$\alpha$+EdU+ OPCs by age. $n_{6\ month\ old}$ = 7, $n_{14\ month\ old}$ = 7, $n_{21\ month\ old}$ = 8. Kruskal-Wallis test, H(2) = 17.25, p=0.0002. Post-hoc comparisons via Steel-Dwass method. Box plots represent median and IQR. **f,** Example image from a *Mobp*-EGFP mouse showing a Bcas1$^{high}$*Mobp*-EGFP- preOL. White dashed circle denotes cell of interest. **g,** Quantification of density of Bcas1$^{high}$*Mobp*-EGFP- preOLs by age. $n_{6\ month\ old}$ = 7, $n_{14\ month\ old}$ = 7, $n_{21\ month\ old}$ = 8. Kruskal-Wallis test, H(2) = 13.39, p=0.0012. Post-hoc comparisons via Steel-Dwass method. Box plots represent median and IQR. **h,** Example image from a *Mobp*-EGFP mouse showing an intact Bcas1$^{high}$*Mobp*-EGFP- preOL and a degenerating Bcas1$^{high}$*Mobp*-EGFP- preOL. Note the fragmenting processes in the degenerating preOL. White dashed circle denotes cell of interest. **i,** Quantification of percentage of Bcas1$^{high}$*Mobp*-EGFP- preOLs that have a degenerating morphology by age. $n_{6\ month\ old}$ = 7, $n_{14\ month\ old}$ = 7, $n_{21\ month\ old}$ = 8. Kruskal-Wallis test, H(2) = 11.95, p=0.0025. Post-hoc comparisons via Steel-Dwass method. Box plots represent median and IQR. **j,** Experimental protocol for assessment of OPC post-division age in young adult and middle-aged mice. Mice were only included in analysis if they were imaged for the maximum duration of 12 weeks ($n_{young\ adult}$ = 3, $n_{middle\text{-}aged}$ = 3). **k**, Percentage of the OPC population that was the indicated post-division age after 12 weeks of imaging. Note the large population of OPCs more than 84 days from cell division in middle-aged animals. Young adult mice are dark grey and middle-aged mice are light blue. $n_{young\ adult}$ = 3, $n_{middle\text{-}aged}$ = 3. Error bars represent S.E.M. Dots denote individual animal means. For all analyses: n.s. not significant, *p < 0.05, **p < 0.01, ***p < 0.001. For detailed statistical information see Supplementary Table 1.

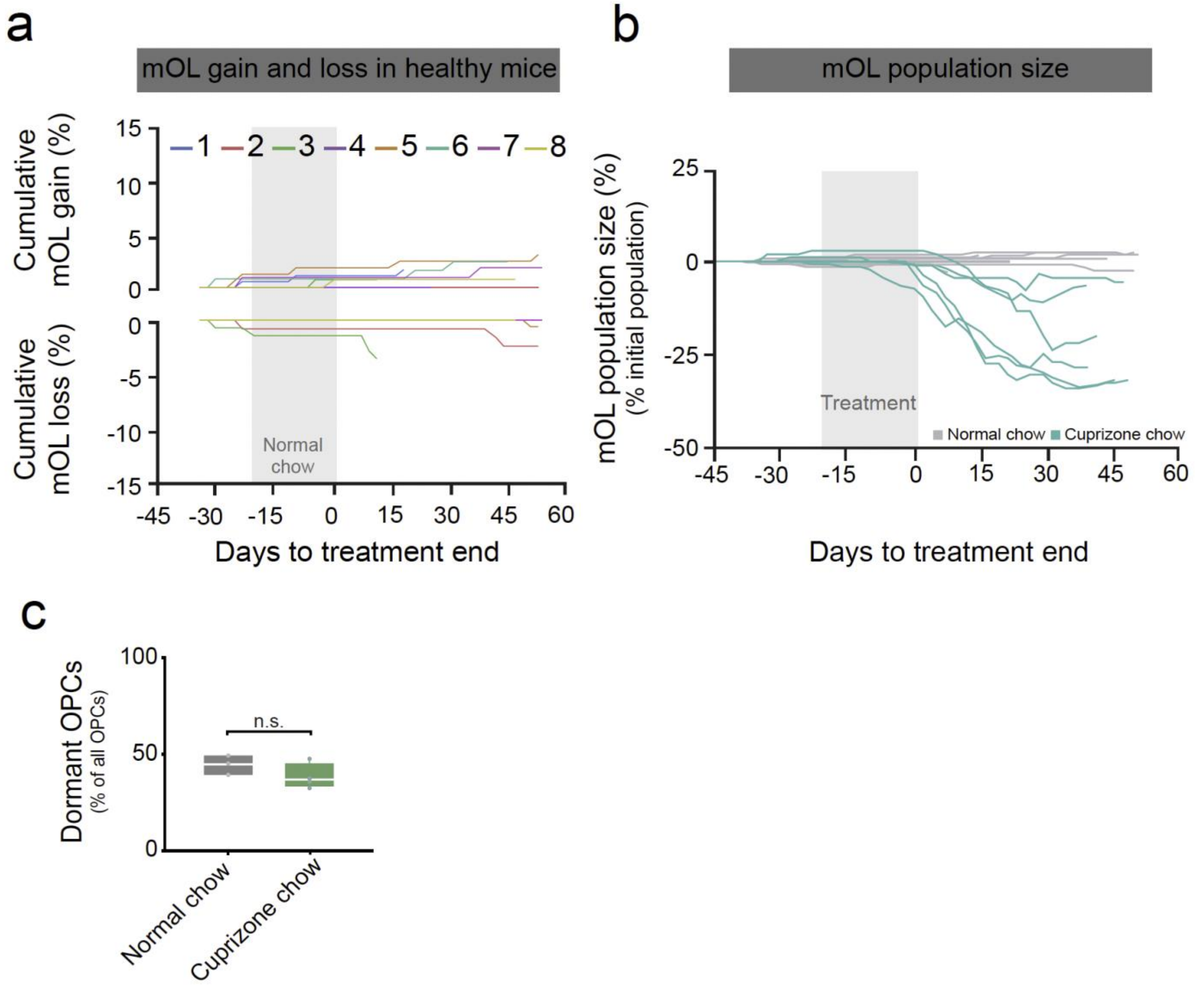


**Supplementary Figure 6**

**Supplementary Figure 6 | Cellular dynamics of oligodendroglia across demyelination and repair in the middle-aged cortex.**

**a,** Cumulative oligodendrocyte gain and loss as a percentage relative to the oligodendrocyte population on day one of imaging in healthy middle-aged mice. The timing of treatment is shown with the grey box. Individual lines represent individual mice ($n_{mice}$ = 8). **b,** The size of the myelinating oligodendrocyte population as a percentage relative to the population on day one of imaging. Lines represent individual mice ($n_{healthy}$ = 8,grey; $n_{cuprizone}$ = 6, green). **c**, Quantification of the percentage of the OPC population that is dormant, having not undergone cell division or differentiation over the course of 12 weeks in healthy and cuprizone fed middle-aged mice. $n_{young\ adult}$ = 3, $n_{middle\text{-}aged}$ = 4. Wilcoxon Exact test, p=0.2286. Dots represent individual mice. Box plots represent median and IQR. For all analyses: n.s. not significant, *p < 0.05, **p < 0.01, ***p < 0.001. For detailed statistical information see Supplementary Table 1.

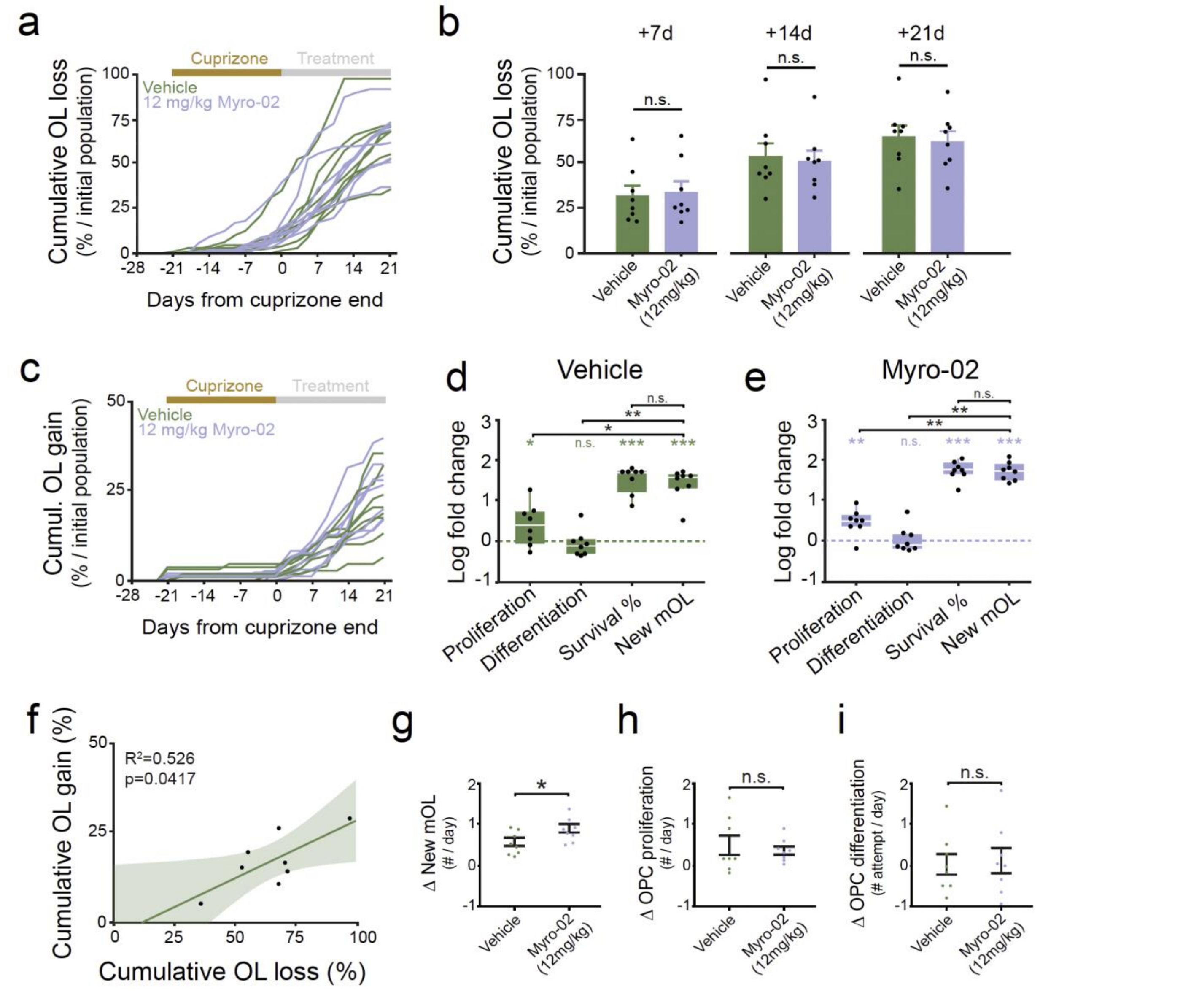


**Supplementary Figure 7**

**Supplementary Figure 7 | Oligodendrocyte gain and loss during Myro-02 treatment.**
**a,** Cumulative oligodendrocyte loss as a percentage of the initial oligodendrocyte population. The timing of treatment is shown with the grey box. Lines represent individual mice **b,** Cross sectional measurements of cumulative oligodendrocyte loss percentage. Day 7: Pooled t-test, t(14) = 0.27, p = 0.7937. Day 14: Pooled t-test, t(14) = 0.28, p = 0.7871. Day 21: Pooled t-test, t(14) = 0.28, p = 0.7803. **c,** Cumulative oligodendrocyte gain as a percentage of the initial oligodendrocyte population. The timing of treatment is shown with the grey box. Lines represent individual mice. **d,** Log fold-change of cell behaviors from baseline to treatment in mice treated with Vehicle. One tailed t-test to determine if log fold-change was greater than zero: Initiation of OPC differentiation (t(7)=-0.06, p=0.6839), OPC proliferation (t(7)=2.24, p=0.0299), preOL survival (t(7)=12.74, p<0.0001), and new mOL (t(7)=10.23, p<0.0001). There was significant effect of cell behavior on log fold-change, Kruskal-Wallis test: H(3) = 22.56, p < 0.0001. Wilcoxon rank sum with Bonferroni correction for post-hoc tests. $n_{mice}$ = 8. **e**, Log fold-change of cell behaviors from baseline to treatment in mice treated with Myro-02. One tailed t-test to determine if log fold-change was greater than zero: Initiation of OPC differentiation (t(7)=0.3177, p=0.3800), OPC proliferation (t(7)=4.16, p=0.0021), preOL survival (t(7)=20.99, p<0.0001), and new mOL (t(7)=21.69, p<0.0001). There was significant effect of cell behavior on log fold-change, Kruskal-Wallis test: H(3) = 24.55, p < 0.0001. Wilcoxon rank sum with Bonferroni correction for post-hoc tests. $n_{mice}$ = 8. **f,** Cumulative oligodendrocyte loss predicts cumulative oligodendrocyte gain on day 21 post-cuprizone in Vehicle treated mice. Linear regression: F(7)=6.67, p = 0.0417, $R^2$=0.526. **g,** Change in the daily number of new oligodendrocytes generated per day from baseline to treatment. Pooled t-test: t(14) = 2.31, p = 0.0365. **h,** Change in the number of OPCs proliferate per day from baseline to treatment. Pooled t-test: t(14) = 0.51, p = 0.6187. **i,** Change in the number of OPCs that attempt differentiation per day from baseline to treatment. Pooled t-test: t(14)

= 0.23, $p = 0.8205$. Error bars denote S.E.M. Dots denote individual animal means. For all analyses: n.s. not significant, *$p < 0.05$, **$p < 0.01$, ***$p < 0.001$. For detailed statistical information see Supplementary Table 1.

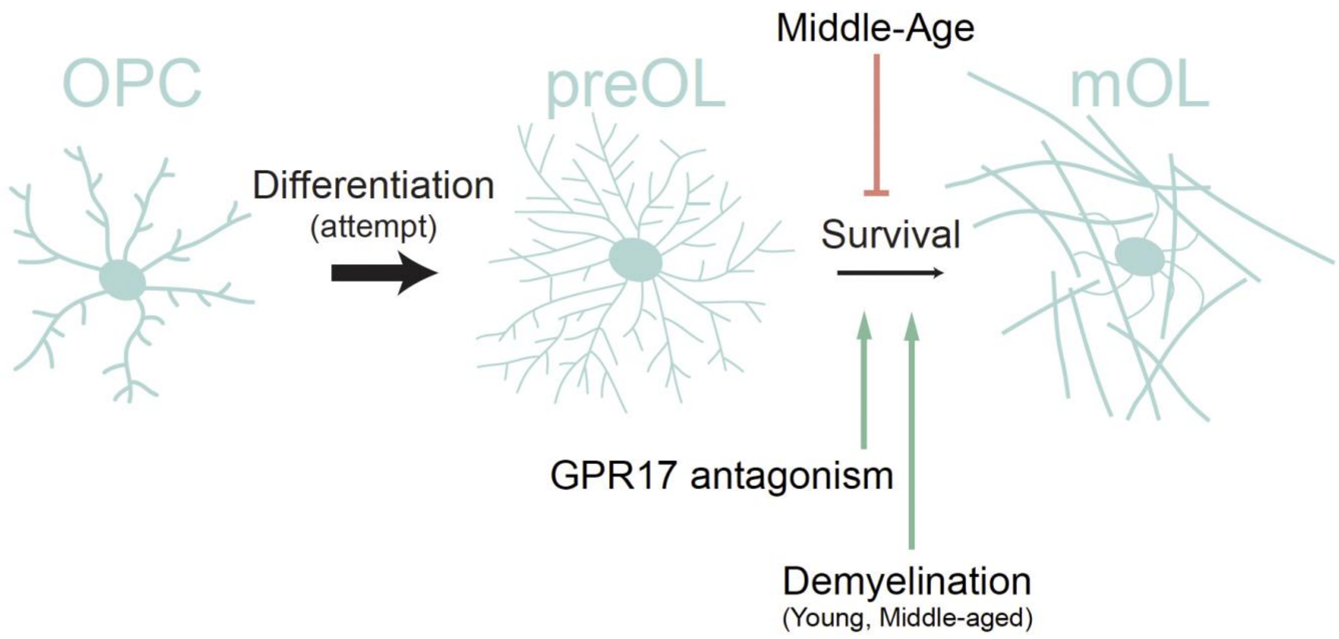


**Supplementary Figure 8**

**Supplementary Figure 8 | Graphical summary: Control of premyelinating oligodendrocyte survival throughout adulthood.**

The production of new myelinating oligodendrocytes varies across health, remyelination and aging**;** however, the cellular mechanisms that this process was not fully understood. Here, we assessed premyelinating oligodendrocyte (preOL) survival during regeneration after demyelination, throughout aging, and during treatment with a GPR17 antagonist. Through these manipulations we provide evidence that preOL survival is modulated throughout adulthood to controls the generation of new myelinating oligodendrocytes.

**GPR17 antagonism:** The GPR17 antagonist, Myro-02, enhances oligodendrocyte replacement after demyelination by promoting the survival of preOLs during differentiation.

**Remyelination:** After cuprizone-mediated demyelination there is an increase in preOL survival and no change in the OPC differentiation. The increase in preOL survival drives the increase in oligodendrocyte generation. This effect was present during regeneration in both young adult and middle-aged mice.

**Aging:** The age-related decline in oligodendrocyte generation is due at least in part to impaired preOL survival. Furthermore, in the middle-aged brain, preOL survival is reduced even within the population of oligodendroglia that exhibit no impairment in cell division or differentiation.
